# Phosphate and phosphite differentially impact the proteome and phosphoproteome of Arabidopsis suspension cell cultures

**DOI:** 10.1101/2020.05.29.124040

**Authors:** Devang Mehta, Mina Ghahremani, Maria Pérez-Fernández, Maryalle Tan, Pascal Schläpfer, William C. Plaxton, R. Glen Uhrig

**Author notes:** **Footnote:** Mina Ghahremani. current address: Department of Biology, University of Ottawa, Ottawa, Ontario, Canada.

## Abstract

Phosphorus absorbed in the form of phosphate (H_2_PO_4_^−^) is an essential but limiting macronutrient for plant growth and agricultural productivity. A comprehensive understanding of how plants respond to phosphate starvation is essential to develop more phosphate-efficient crops. Here we employed label-free proteomics and phosphoproteomics to quantify protein-level responses to 48 h of phosphate versus phosphite (H_2_PO_3_^−^) resupply to phosphate-deprived *Arabidopsis thaliana* suspension cells. Phosphite is similarly sensed, taken up, and transported by plant cells as phosphate, but cannot be metabolized or used as a nutrient. Phosphite is thus a useful tool to delineate between non-specific processes related to phosphate sensing and transport, and specific responses to phosphorus nutrition. We found that responses to phosphate versus phosphite resupply occurred mainly at the level of protein phosphorylation, complemented by limited changes in protein abundance, primarily in protein translation, phosphate transport and scavenging, and central metabolism proteins. Altered phosphorylation of proteins involved in core processes such as translation, RNA splicing, and kinase signalling were especially important. We also found differential phosphorylation in response to phosphate and phosphite in 69 proteins, including splicing factors, translation factors, the PHT1;4 phosphate transporter and the HAT1 histone acetyltransferase—potential phospho-switches signalling changes in phosphorus nutrition. Our study illuminates several new aspects of the phosphate-starvation response and identifies important targets for further investigation and potential crop improvement.

## INTRODUCTION

Phosphorus (P) is an essential macronutrient for plant growth that is absorbed by roots in its fully oxidized and anionic form, orthophosphate (H_2_PO_4_^−^; Pi). Pi is an essential structural constituent of important biomolecules such as nucleic acids, phospholipids, and sugar-phosphates. Pi also plays central roles in photosynthesis and respiration, as well as in signal transduction and metabolic regulation via its covalent attachment to phosphoproteins. Despite its importance, soluble Pi is notably scarce as a bioavailable macronutrient in most soils globally and is therefore widely used in fertilizer formulations. However, most crops only assimilate 20-30% of applied Pi, with the rest either being precipitated in the soil as insoluble metal cation–Pi complexes, converted into organic-P compounds by soil microbes, or leached into water bodies causing algal blooms and eutrophication (Lambers and Plaxton, 2015). Furthermore, Pi fertilizers are manufactured from rock–Pi, an unsustainable resource predicted to be depleted globally within the next 100 years (Cordell and White, 2014). Hence, it is vital to gain a better understanding of how plant cells sense and respond to changes in Pi availability in order to identify potential targets for breeding and biotechnological interventions that will produce the Pi-efficient crop varieties urgently needed for long-term global food security and ecosystem preservation.

Intracellularly, phosphorus is present in plant cells both as free Pi as well as organic phosphate-esters such as nucleotides, phospholipids, and phosphoproteins, of which the latter, results from the trans-phosphorylation of target proteins by protein kinases. Protein kinase-mediated protein phosphorylation is the most abundant post-translational modification (PTM) across all eukaryotes, representing a primary means for cellular signalling and metabolic control (Jin and Pawson, 2012). At least 75% of all eukaryotic proteins appear to be phosphorylated throughout their lifetime (Sharma et al., 2014). In plants, multiple studies have indicated extensive protein phosphorylation across multiple tissues, organs and developmental stages (Uhrig et al., 2019; Mergner et al., 2020) as well as in response to nutrient stress (Engelsberger and Schulze, 2012; Duan et al., 2013; Li et al., 2014; Menz et al., 2016; Yang et al., 2019; Saito and Uozumi, 2020). As protein phosphorylation is integral to plant cell functionality, deciphering the dynamics of protein phosphorylation under Pi-limiting versus Pi-replete conditions is essential to understanding how gene expression and metabolism are controlled during plant acclimation to Pi-deficient soils, in addition to the key signalling events that convey changes in Pi nutritional status. Furthermore, due to the dependence of protein kinase signalling on Pi through ATP availability, studying how Pi nutrition impacts protein phosphorylation could potentially delineate essential versus non-essential signalling events when plants respond to nutrient stress.

In order to mitigate the deleterious consequences of Pi deprivation, Pi-starved (−Pi) plants exploit complex local and systemic signaling pathways that elicit an array of morphological, physiological, and metabolic adaptations known as the Pi-starvation response (PSR). Although these adaptations are by no means identical in all plants, certain aspects are conserved in a broad variety of species from diverse environments. The PSR arises from coordinated changes in gene and protein expression, along with various post-translational controls, that lead to reprioritization of internal Pi use while enhancing external Pi acquisition (Zhang et al., 2014). For example, Pi deficiency results in altered root phenotypes (e.g. increased root hair density and root-to-shoot ratios) and augmented metabolism that drives the release of carboxylates and Pi scavenging enzymes into the rhizosphere to enhance Pi availability for uptake (Lambers and Plaxton, 2015).

A combination of systemic ‘omic’ (Li et al., 2008; Tran and Plaxton, 2008; Plaxton and Tran, 2011) and targeted biochemical (Gregory et al., 2009; Liang et al., 2012; Shane et al., 2013; Ghahremani et al., 2019) studies have indicated that multiple PTMs play important roles in the PSR. These PTMs include phosphorylation, ubiquitination, and glycosylation signatures that impact gene expression profiles and the activities of intra- and extracellular metabolic enzymes (Shane et al., 2013; Plaxton and Shane, 2015; Ghahremani et al., 2019; Pan et al., 2019). Targeted studies involving protein phosphorylation indicate that calcium dependent protein kinases (CDPKs) are likely protein-level PTM regulators of the PSR (Saito and Uozumi, 2020), which aligns with observed fluctuations in calcium levels under changing Pi conditions (Matthus et al., 2019). Despite these advancements, much remains to be resolved about the PSR, including elucidation of key proteins involved in Pi signaling in addition to determining the corresponding protein target landscape of these signalling molecules. Our study combines quantitative proteomics and phosphoproteomics to help answer these questions, while advancing potential solutions urgently needed to optimize Pi-fertilizer inputs in agriculture by identifying novel protein targets of potential biotechnological use (Veneklaas et al., 2012).

To identify alternative sources for phosphorus nutrition in plants, researchers have previously attempted to repurpose a related chemical, phosphite (H_2_PO_3_^−^; Phi, also known as phosphonate) as a combined fertilizer and fungicide (López-Arredondo and Herrera-Estrella, 2012; Jost et al., 2015; Lambers and Plaxton, 2015). Phi is a reduced form of Pi, in which a hydrogen replaces an oxygen bonded to the P atom, and is widely used as a systemic, phloem-mobile fungicide to control crop diseases caused by various oomycete species belonging to the genus Phytophthora (McDonald et al., 2001; Lambers and Plaxton, 2015). Although Phi-treated plants rapidly transport and concentrate Phi (using Pi-transporters), it is metabolically inert in plant cells and thus persists in their tissues for extensive periods (McDonald et al., 2001). Phi does exert direct effects on plants, being extremely phytotoxic to −Pi, but not Pi-sufficient plants (Carswell et al., 1996; Carswell et al., 1997; McDonald et al., 2001; Ticconi et al., 2001; Singh et al., 2003; Lambers and Plaxton, 2015). This is because Phi blocks the acclimation of plants (and yeast) to Pi deficiency by disrupting the de-repression of genes encoding proteins characteristic of their PSR (e.g., repressible acid phosphatases [APase] and high-affinity plasmalemma Pi transporters, etc.) (Carswell et al., 1996; Carswell et al., 1997; McDonald et al., 2001; Ticconi et al., 2001; Karthikeyan et al., 2002). The Phi anion therefore represents a valuable tool to manipulate and investigate how plants monitor, signal, and hence generate an integrated response to Pi deprivation.

The aim of the present study was to exploit high-resolution liquid chromatography-tandem mass spectrometry (LC-MS/MS) to obtain a comprehensive overview of the impact of Pi versus Phi resupply on the proteome and phosphoproteome of heterotrophic suspension cell cultures of the model plant *Arabidopsis thaliana*. This approach allowed us to elucidate cellular responses that are uniquely reliant on Pi-metabolism rather than the activation of common Pi / Phi sensing mechanisms. Uniquely, our study also helps to better define the molecular mechanisms by which Phi induces toxicity to −Pi plant cells, which holds promise for potential agricultural biotechnology benefits.

## RESULTS AND DISCUSSION

In order to perform the starvation and re-feeding experiment, we initially cultured Arabidopsis suspension cells for 6 days in Murashige-Skoog media containing 5 mM KPi, a level needed to maintain these cells fully Pi-sufficient over 6 days in batch culture (Veljanovski et al., 2006; Tran et al., 2010b). The cells were then subcultured into Pi-free media for 3 days, after which each −Pi flask was directly supplemented with either no Pi, 2 mM Pi (+Pi), or 2 mM Phi (+Phi) and cultured for an additional 2 days prior to harvest (Figure 1A). To validate the experimental samples we first assayed APase activity in clarified extracts, whose upregulation is a well-known marker of the plant PSR (Veljanovski et al., 2006). As expected, 48 h of Pi or Phi resupply elicited a dramatic (~4-fold) reduction in the extractable APase activity of the −Pi cells (Figure 1B). Further sample validation employed immunoblotting using phosphorylation site-specific antibodies (anti-pSer11) to monitor the phosphorylation status of phosphoenolpyruvate carboxylase (PEPC) (Tripodi et al., 2005). PEPC is a tightly regulated enzyme situated at a key branch point of cytosolic glycolysis, and that plays important roles in the metabolic adaptations of −Pi plants (Gregory et al., 2009; O’Leary et al., 2011b; Plaxton and Tran, 2011; Shane et al., 2013). The anti-pSer11 antibodies detect the well-characterized and highly conserved N-terminal seryl phosphorylation site of plant-type PEPCs (PTPC), which when present is a specific indicator of PTPC activation (O’Leary et al., 2011a). Plant Pi deprivation prompts enhanced PTPC phosphorylation and activity, as well as a 300% increase in *in vivo* flux through the PEPC reaction (Masakapalli et al., 2014), along with increased expression of malate dehydrogenase, citrate synthase, and plasma membrane-targeted carboxylate transporters (Gregory et al., 2009; Plaxton and Tran, 2011). The immunoblots of Figure 1C demonstrated that Pi or Phi resupply to the −Pi cells elicited *in vivo* dephosphorylation of 107 kDa PTPC subunits at their regulatory N-terminal serine phosphosite. This was corroborated following mass spectrometry analysis (LC-MS/MS) of TiO_2_-enriched phosphopeptides derived from the respective proteomes in which the PTPC isozymes AtPPC1 and AtPPC3 were detected (Figure 2B, Tables S1 and S2). Taken together, we have successfully created Pi-starvation conditions, and recovery from Pi-deprivation through Pi resupply.

**Figure 1.**
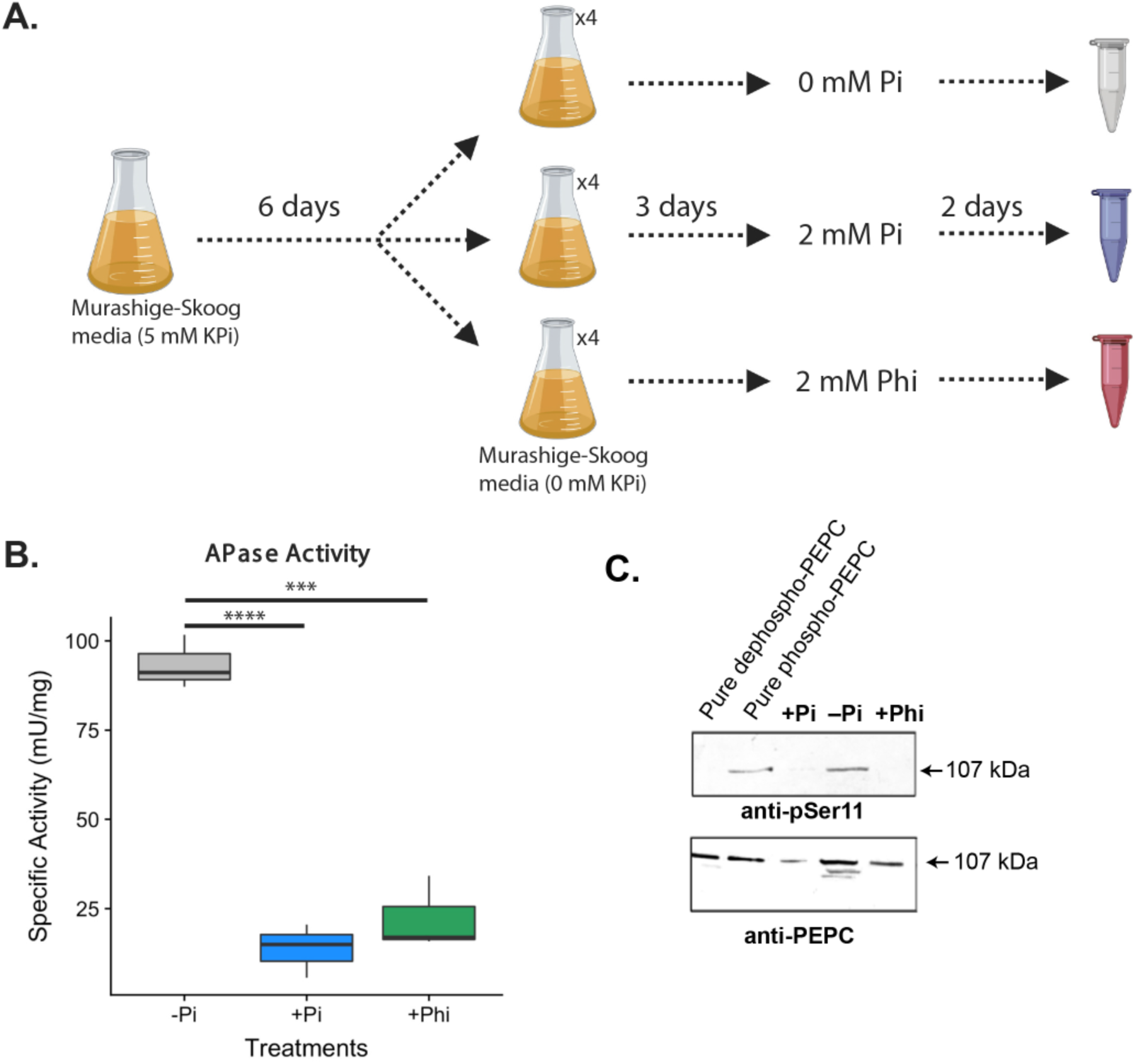
Sample preparation and validation. (A) Experimental workflow for preparation of −Pi, +Pi, and +Phi Arabidopsis suspension cells. (B) APase specific activities of the respective extracts represent means ±SE of *n* = 3 biological replicates. Significant differences were determined using the Student’s *t*-test; *p value* <0.05 (***), *p value* <0.01 (****). (C) Clarified extracts from the −Pi, +Pi and +Phi cells were subjected to SDS/PAGE (12 μg protein/lane) and electroblotted on to a PVDF membrane. Immunoblots were probed with either a phosphosite-specific antibody raised against the conserved N-terminal Ser11 phosphorylation domain of a castor bean PEPC isozyme (anti-pSer11) (Tripodi et al., 2006), or a PEPC-specific antibody raised against the corresponding, fully purified castor bean PEPC (anti-PEPC) (Gennidakis et al., 2011). ‘Dephospho-PEPC’ and ‘Phospho-PEPC’ correspond to 50 ng each of the PEPC isozyme AtPPC1 that had been fully purified from the −Pi Arabidopsis suspension cells and preincubated with and without λ protein phosphatase, respectively, as described by Gregory and co-workers (2009).

**Figure 2.**
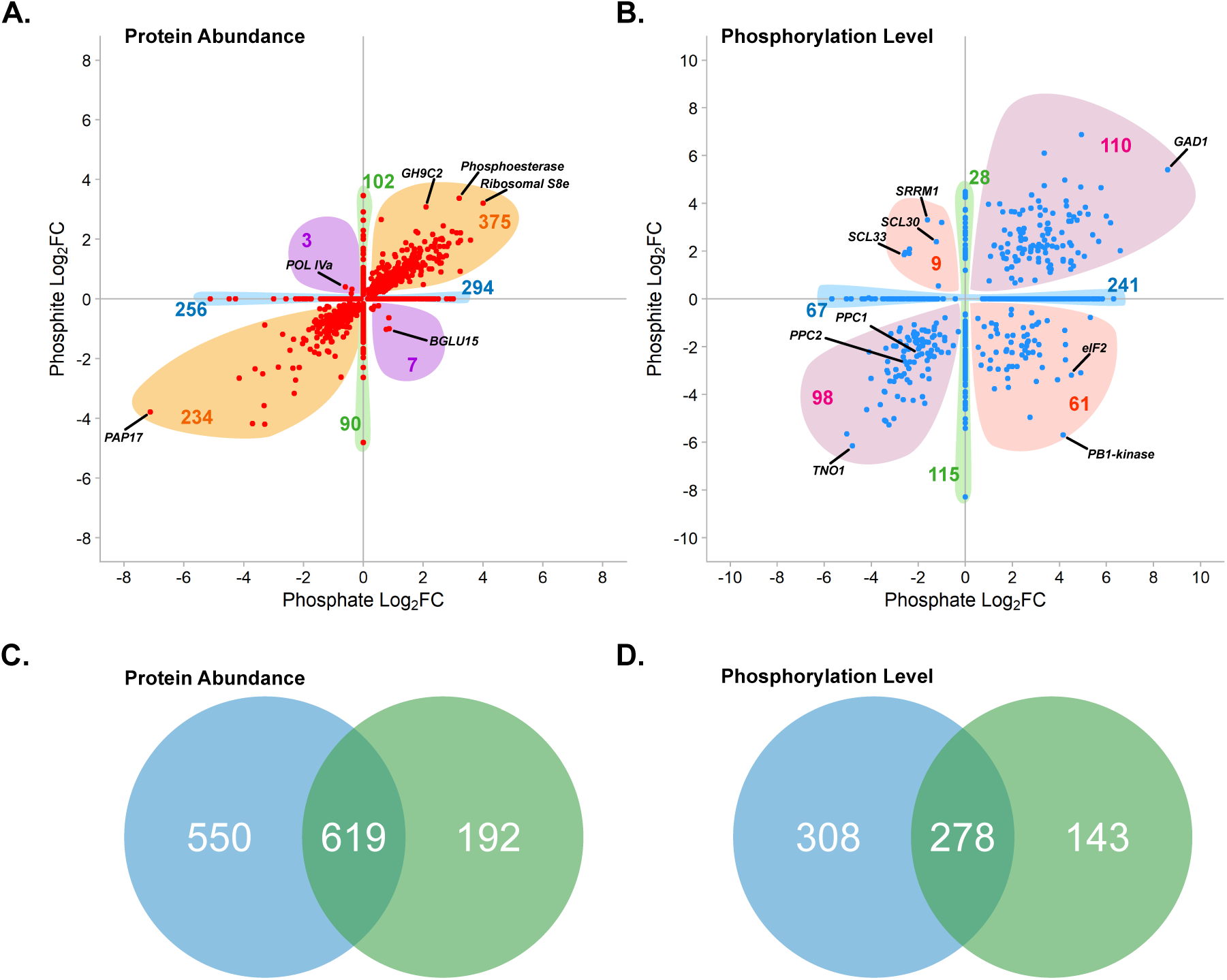
Distribution of significant proteome and phosphoproteome fluctuations in response to re-feeding with Phi or Pi. **(A-B)** Distribution of significantly changing proteome and phosphoproteome (q-value <0.05), respectively (n=4). Numbers denote the number of significantly changed species on each axis or in each quadrant. Labeled proteins represent those with the largest Log_2_FC in total abundance or phosphorylation status. **(C-D)** Venn diagrams depicting the overlap of significantly changing proteins or phosphorylation events common between or independent of the Phi and Pi treatments (q-value<0.05; n=4).

We next exploited high-resolution LC-MS/MS to quantitatively assess +Pi and +Phi induced proteome and phosphoproteome changes relative to −Pi. Our analysis identified nearly 5,100 proteins and 1,600 phosphoproteins, quantifying changes in the abundance of 3,270 proteins and the phosphorylation status of 1,361 phosphoproteins respectively, across all three experimental conditions (Table 1). Of these quantified proteins and phosphoproteins, 1,114 (+Pi vs −Pi), 767 (+Phi vs −Pi) and 578 (+Pi vs +Phi) proteins exhibited significant changes in protein abundance *(q-value* ≤ *0.05)*, while 587 (+Pi vs −Pi), 432 (+Phi vs −Pi) and 552 (+Pi vs +Phi) phosphoproteins exhibited significant changes in their phosphorylation status *(q-value* ≤ *0.05)* (Table 1). We next queried phosphoproteins with significantly changing phosphorylation sites against the significantly changing proteome. This revealed that the majority of significantly changing phosphorylated proteins do not exhibit a significant change in abundance, suggesting that the observed phosphorylation changes are predominantly due to fluctuations in phosphorylation status and not changes in abundance (Table 2). Together, our acquisition of both proteomic and phosphoproteomic data provides a comprehensive, single study understanding of the global protein-level changes induced by Pi and Phi in Arabidopsis.

**Table 1.**
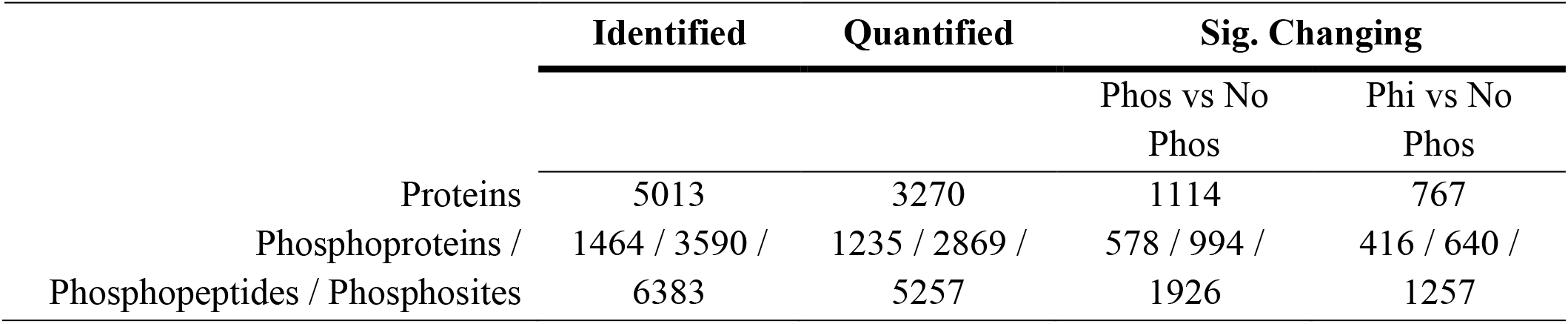
Number of identified, quantified and significantly changing proteins (*q-value* ≤ 0.05), phosphopeptides, and phosphoproteins from −Pi Arabidopsis cell cultures that were resupplied with either 2 mM Pi or Phi for 48 h.

**Table 2.** Comparison of proteins with significantly changing phosphorylation events (Sig. P) to the quantified total proteome (All TP) and significantly changing total proteome (Sig. TP). Significantly changing species possessed a *q-value* ≤ *0.05*.

|  | Phosphite (Phi) |  | Phosphate (Pi) |  |
| --- | --- | --- | --- | --- |
|  | Sig. P vs All TP | Sig. P vs Sig TP | Sig. P vs All TP | Sig. P vs Sig TP |
| Total number of<br>All +P proteins | 44 | 17 | 147 | 84 |
| % of All +P<br>proteins | 4.6 | 1.8 | 9.7 | 5.5 |
| % of All Sig +P<br>proteins | 72.1 | 27.9 | 63.6 | 36.4 |

### Responses to Pi and Phi occur via changes in protein phosphorylation and not protein abundance

A total of 1,169 and 811 proteins changed significantly in abundance *(q-value* ≤ *0.05)*) upon Pi and Phi resupply, respectively. A total of 619 significantly changing proteins were common to both Pi and Phi re-feeding responses (Table S1). We then plotted the log-transformed fold-changes of proteins significantly changing in +Pi and +Phi conditions (Figure 2A). Here, we also observe a high degree of concordance in protein abundance changes between the two conditions, with only 10 proteins having opposing changes in abundance between the +Pi and +Phi samples (7 proteins were upregulated under +Pi and downregulated under +Phi while only 3 proteins were downregulated under +Pi and upregulated under +Phi conditions; Figure 2B). Conversely, a total of 609 proteins changed similarly in +Pi and +Phi conditions. Overall the degree of change in protein abundance was greater under +Pi conditions than under +Phi, compared to −Pi. This indicates that proteome-level changes are more sensitive to Pi than to Phi. Overall, these results corroborate previous findings demonstrating that +Phi attenuates the plant PSR (Ticconi et al., 2001), albeit to a lower degree than +Pi.

Quantitative assessment of the phosphoproteome found 556 proteins significantly changing in their phosphorylation status upon +Pi and 421 proteins upon +Phi. Among these, 278 phosphoproteins changed significantly in both +Pi and +Phi conditions (Figure 2C). However, unlike changes in protein abundance, we observed limited concordance in the degree of phosphorylation change between +Pi and +Phi conditions (Figure 2B). Out of the 278 commonly changing phosphoproteins, 70 exhibited differential changes between the two conditions. (61 proteins showed increased phosphorylation under +Pi and decreased phosphorylation under +Phi; 9 proteins showed increased phosphorylation under +Phi and decreased phosphorylation under +Pi; Figure 2B). We then assessed the relative contributions of each subcellular compartment to changes in protein abundance and protein phosphorylation under +Pi and +Phi using the SUBA database (SUBA4; http://suba.live/; (Hooper et al., 2017). An increase in the proportion of significantly changing proteins from the nucleus and plastid along with a corresponding decrease in nuclear and plasma-membrane localised phosphoproteins occurred upon +Pi relative to +Phi (Figure S1).

### Pi and Phi similarly affect the abundance of Pi-sensing, transport, and scavenging proteins

Plants respond to Pi starvation by upregulating high affinity inter- and intracellular Pi transporters, and intra- and extracellular Pi scavenging enzymes (e.g. primarily purple acid phosphatases (PAPs); (Tran et al., 2010a)). Five families of Pi transporters (PHT1-4 and PHO1) have been identified in plants, each responsible for Pi transport across different cellular or intracellular membranes (Rouached et al., 2010; Versaw and Garcia, 2017). Of these, the PHT1 family of high affinity H_2_PO_4_^−^ /H^+^ symporters has 9 plasma membrane localized members in Arabidopsis. In our data we observe significant decreases in protein abundance for PHT1;2 (AT5G43370; +Pi: −2.69 Log_2_FC; +Phi: −1.18 Log_2_FC), PHT1;4 (AT2G38940; +Pi: −3.29 Log_2_FC; +Phi: −0.87 Log_2_FC) and PHT1;6 (AT5G43340; +Pi: −4.15 Log_2_FC; +Phi: −2.65 Log_2_FC) in response to both +Pi and +Phi. This observation provides protein-level support to previous studies showing repression of *PHT1 transporter* expression in +Phi conditions and the ability of high affinity Pi transporters to also transport Phi (Ticconi et al., 2001; Karthikeyan et al., 2002). High affinity PHT1 transporters in Arabidopsis are localized to the plasma membrane through the action of PHOSPHATE TRANSPORTER TRAFFIC FACILITATOR1 (PHF1; AT3G52190), a plant-specific Sec-12 related protein present in the endoplasmic reticulum (González et al., 2005). We also detected significant decreases in PHF1 protein abundance in both +Pi and +Phi conditions (+Pi: −1.2 Log_2_FC; +Phi: −0.25 Log_2_FC), further supporting a non-specific Pi/Phi response by PHT1 transporter expression.

Pi-deprived plants also upregulate a suite of hydrolytic enzymes to scavenge Pi from extra- and intracellular Pi esters (Tran et al., 2010a). These include lipases that facilitate the plastidial production of galacto- and sulfolipids for cell-wide replacement of membrane phospholipids (Tran et al., 2010a). Correspondingly, we observed significant downregulation of NON-SPECIFIC PHOSPHOLIPASE C3 (NPC3; AT3G03520) (+Pi: −0.78 Log_2_FC; +Phi: −0.66 Log_2_FC) and PHOSPHOLIPASE 2 (PLC2; AT3G08510) (+Pi: −2.33 Log_2_FC; +Phi: −1.35 Log_2_FC) in both +Pi and +Phi conditions. NPC3 is a member of the Group 2 NPC proteins that have well-characterized roles in the replacement of phospholipids by galacto- and sulfo-lipids during Pi starvation in plants (Nakamura et al., 2005; Gaude et al., 2008). An Arabidopsis T-DNA knockout mutant of NPC3 displayed decreased lateral root densities (Wimalasekera et al., 2010), further supporting its role in Pi starvation. PLC2 is a phosphoinositide-specific phospholipase that has not been studied in the context of plant Pi starvation. However, the strong decrease in PLC2 protein abundance in both +Pi and −Pi conditions (+Pi: −2.33 Log_2_FC; +Phi: −1.35 Log_2_FC), coupled with prior studies showing the importance of other inositol phosphate metabolic enzymes in maintaining Pi homeostasis (Kuo et al., 2018), suggest possible roles for PLC2 in Pi starvation. We also observed +Pi-specific regulation of additional two phospholipases, PHOSPHOLIPASE Dζ2 (PLDζ2; AT3G05630) and PHOSPHOLIPASE Dδ (PLDδ; AT4G35790). We find that PLDζ2 protein abundance decreases significantly only in response to +Pi re-feeding in Arabidopsis cells (−2.15 Log_2_FC). This protein-level data echoes previous findings that PLDζ2 transcript accumulation is increased only in response to Pi and not Phi (Jost et al., 2015) and that PLDζ2 mutant plants have an impaired ability to replace phospholipids with galactolipids upon Pi-starvation (Su et al., 2018). We find that PLDδ protein abundance significantly increases (0.25 Log_2_FC) in response to +Pi and is insensitive to +Phi. PLDδ is known to function in phosphatidic acid biosynthesis and knockout mutants are more sensitive to peroxide-mediated cell death (Zhang et al., 2003). While PLDδ has not been studied in the context of Pi nutrition, the Pi-specific reduction in PLDδ levels might indicate a role in Pi-starvation related cell-death responses.

The replacement of phospholipids by sulfolipids requires the biosynthesis of new sulfoquinovosyldiacylglycerol. In Arabidopsis this reaction proceeds in two steps: the synthesis of UDP-sulfoquinovose from UDP-glucose and sulfite followed by the transfer of the sulfoquinovose moiety from UDP-sulfoquinovose to diacylglycerol (Yu et al., 2002). SULFOQUINOVOSYLDIACYLGLYCEROL 1 (SQD1; AT4G33030) catalyzes the first step of this reaction in response to Pi starvation (Essigmann et al., 1998) and in our study is significantly reduced in abundance upon both +Pi and +Phi (+Pi: −1.44 Log_2_FC; +Phi: −0.32 Log_2_FC). SULFOQUINOVOSYLDIACYLGLYCEROL 2 (SQD2; AT5G01220) is a sulfoquinovose transferase that catalyzes the second reaction in sulfoquinovosyldiacylglycerol biosynthesis and mutant alleles of SQD2 have previously been shown to have impaired growth under Pi-limiting conditions (Yu et al., 2002). Here, we only observed changes in SQD2 protein abundance under +Pi (−1.9 Log_2_FC). More recently, researchers have found that SQD2 but not SQD1 plays a role in the biosynthesis of a newly discovered plant lipid, glucuronosyl diacylglycerol that protects plants under Pi starvation (Okazaki et al., 2013). Our findings indicate that further investigation is needed to study if the biosynthesis of glucuronosyl diacylglycerol, unlike sulfoquinovosyldiacylglycerol, is specific to Pi nutrition.

A well-characterized aspect of the plant PSR is the induction of specific vacuolar and secreted purple APase (PAP) isozymes (Tran et al., 2010a). The Arabidopsis genome encodes 29 different PAPs with differing subcellular localizations, organ-specific expression, and in Pi metabolism. Here, we observed a significant decrease in the protein abundance of PAP1 (AT1G13750; +Pi: −1.19 Log_2_FC; +Phi: −1.07 Log_2_FC), PAP14 (AT2G46880, +Pi: −2.19 Log_2_FC; +Phi: −2.13 Log_2_FC), PAP17 (AT3G17790; +Pi: −7.12 Log_2_FC; +Phi: −3.78 Log_2_FC), PAP24 (AT4G24890; +Pi: −1.29 Log_2_FC; +Phi: −0.34 Log_2_FC) and PAP25 (AT4G36350; +Pi: −1.4 Log_2_FC; +Phi: −0.59 Log_2_FC) in both the +Pi and +Phi conditions (compared to −Pi samples). Interestingly, we also detected significant decreases in the abundance of two additional Pi-starvation inducible PAPs, AtPAP10 (AT2G16430; +Pi: −0.6 Log_2_FC) and AtPAP12 (AT2G27190; +Pi: −1.32 Log_2_FC) only in response to +Pi, but not +Phi, suggesting possible specificity in their regulation compared to other PAPs detected in our experiment. PAP10, PAP12, PAP17, and PAP25 have been biochemically and functionally characterized in −Pi Arabidopsis, and are strongly induced at transcript and protein level upon Pi-starvation, but rapidly repressed upon +Pi (Tran et al., 2010b; Lan et al., 2012; Del Vecchio et al., 2014; Wang et al., 2014). Interestingly, PAP17 stands out as being the most downregulated of all proteins detected in the +Pi cells, relative to −Pi controls (Fig. 2A; Table S1). PAP17 also exhibits peroxidase activity suggesting a possible role in general stress responses in addition to Pi starvation (Tran et al., 2010a).

### Bypass enzymes of central carbon metabolism and respiration are downregulated by both Pi and Phi

During prolonged Pi stress, depletion of vacuolar Pi pools leads to a marked reduction in the cytoplasmic concentrations of Pi and organic-P metabolites, including nucleotide-Ps such as ATP and ADP (Plaxton and Tran, 2011). This decline is believed to restrict the activity several enzymes of classical glycolysis and mitochondrial respiration that are dependent on ATP, ADP and/or Pi as co-substrates. However, in contrast to Pi and the adenylates, the cytosolic level of inorganic pyrophosphate (PPi) in −Pi plants remains remarkably stable (Plaxton and Tran, 2011). Evidence that −Pi plants favor the use of PPi-dependent cytosolic bypass enzymes is also evidenced by their upregulation of the PPi-dependent proton pump of the tonoplast, sucrose synthase and UDP-glucose pyrophosphorylase, PPi-dependent phosphofructokinase (PFP), pyruvate Pi dikinase (PPDK), and PEPC (Nasr Esfahani et al., 2016). Here, we observed marked decreases in the abundance of several of these bypass enzymes under both +Pi and +Phi treatments (Supplemental Table S1).

Plant cytosolic glycolysis begins with the synthesis of hexose-phosphates from sucrose. This occurs through the hydrolysis of sucrose to fructose and glucose followed by the action of hexokinase and fructokinase to produce glucose-6-phosphate and fructose-6-phosphate; this is energetically expensive, requiring 2 ATP molecules. However, an alternative pathway exists in plants in which sucrose synthase uses sucrose and UDP to produce fructose and UDP-glucose. Fructose is then phosphorylated to fructose-6-phosphate by fructokinase using 1 ATP, and UDP-glucose converted to glucose-1-phosphate by UDP-glucose pyrophosphorylase. The latter reaction uses PPi and produces UTP as a byproduct and is thus likely preferred during Pi starvation. In support of this hypothesis, we find that sucrose synthase-4 (SUS4; AT3G43190; +Pi: −1.14 Log_2_FC; +Phi: −0.53 Log_2_FC) protein abundance was reduced in both +Pi and +Phi cells. In addition, sucrose synthase-1 and -3 levels were also reduced in +Pi samples. Furthermore, the protein abundance of UDP-glucose pyrophosphorylase-1 (UGPase; AT3G03250; +Pi: −0.22 Log_2_FC; +Phi: −0.31 Log_2_FC) was also reduced in both +Pi and +Phi cells.

The oxidative pentose-phosphate pathway (OPPP) is an alternative route of hexose-phosphate metabolism that produces reducing power (i.e. NADPH) and carbon skeletons (e.g. ribose-5-phosphate, erythrose-4-phosphate) needed for various biosynthetic pathways. Interestingly, glucose-6-phosphate dehydrogenase-6 (G6PDH-6; AT5G40760.2), which catalyzes the first committed step in the OPPP, was one of the most significantly phosphorylated metabolic enzymes in both +Pi and +Phi cells; G6PDH-6 was bisphosphorylated at high stoichiometry at adjacent seryl and threonine residues located near its N-terminus (Supplemental Table S2). G6PDH-6 is the dominant G6PDH isozyme expressed throughout Arabidopsis plants (Wakao and Benning, 2005). To best of our knowledge, however, there have been no reports regarding the functions or mechanisms of plant G6PDH phosphorylation. It will therefore be of considerable interest to test the hypothesis that phosphorylation activates G6PDH-6 for enhancing the flux of hexose-phosphates into the OPPP, in support of the rapid resumption of anabolism and overall plant growth that ensues Pi resupply to −Pi plants (Plaxton and Tran, 2011)

In the first committed step of glycolysis, fructose-6-phosphate is converted into fructose-1,6-bisphosphate. This reaction is typically catalyzed by ATP-dependent phosphofructokinase (PFK). However, under ATP limited conditions, such as during Pi-deprivation or anoxia, this reaction can also be carried out by PFP in plants. We observed a significant reduction in PFP β-subunit protein abundance (AtPFPβ2; AT4G04040; +Pi: −1.63 Log_2_FC; +Phi: −0.75 Log_2_FC) under both +Pi and +Phi conditions. While AtPFPβ2 has not specifically been studied under −Pi conditions, AtPFPβ2 RNAi knockdown lines exhibited growth retardation and reduced CO_2_-assimilation, under normal growth conditions (Lim et al., 2009).

The final reaction in classical glycolysis involves the irreversible conversion of phosphoenolpyruvate (PEP) and ADP into pyruvate and ATP. Plants possess several bypasses for this step that avoid the use of ADP. This includes a series of reactions beginning with the irreversible β-carboxylation of PEP to oxaloacetate, catalyzed by PEPC. This oxaloacetate is reduced into malate by cytosolic NAD-malate dehydrogenase (c-NAD-MDH) and malate is then transported into the mitochondria where it is oxidized into pyruvate by NAD-malic enzyme. We observed a downregulation of abundance of the PEPC isozymes PPC1 and PPC3 in +Pi and +Phi cells (Supplemental Table S1). PEPC activity was also regulated through phosphorylation during +Pi and +Phi as discussed above (Fig. 1C) and we also observed striking decreases in phosphorylation of PPC1 and PPC3 during both +Pi and +Phi (PPC1: AT1G53310; +Pi: −2.121 Log_2_FC; +Phi: −1.78 Log_2_FC and PPC3: AT2G42600; +Pi: −2.63 Log_2_FC; +Phi: −2.42 Log_2_FC). We also detected a decrease in the abundance of c-NAD-MDH1 only in +Pi cells (AT1G04410; +Pi: −0.63).

Interestingly, one of the most phosphorylated proteins in our entire dataset was glutamate decarboxylase-1 (GAD1; AT5G17330; +Pi: 8.60 median Log_2_FC; +Phi: 5.41 median Log_2_FC), with multiple N-terminal seryl phosphorylation sites increasing in response to +Pi and +Phi (Supplemental Table S2). GAD exists in both plants and animals, catalyzing the irreversible decarboxylation of glutamate into γ-aminobutyric acid (GABA) in the cytosol as the first committed step of a short metabolic pathway called the GABA shunt. The GABA shunt bypasses the first reactions of the mitochondrial TCA cycle to produce succinate, which ultimately helps drive ATP production through mitochondrial respiration under multiple stresses (Michaeli and Fromm, 2015). The role(s) of these multiple, closely spaced, and highly conserved N-terminal GAD phosphorylation sites in plants remains uncharacterized; however, in the human ortholog HsGAD2, phosphorylation at equivalent sites triggers HsGAD2’s association with the membranes of neurotransmitter (i.e. GABA) releasing vesicles (Namchuk et al., 1997). Despite a clear role(s) for GABA in plants remaining largely elusive (Bown and Shelp, 2020), our findings, coupled with the phosphorylation mediated vesicle association of HsGAD2, suggests that GABA metabolism in plants may take on Pi nutrition dependent roles that are dictated by phosphorylation-mediated changes in GAD’s subcellular localization (e.g. strictly cytosolic versus membrane associating).

Lastly, the mitochondrial electron transport chain is heavily regulated by the availability of ADP and Pi. Under ADP and Pi limited conditions plant mitochondria can utilise the non-proton motive alternative oxidase (AOX) to decouple electron transfer from ATP synthesis, thereby bypassing the activities of Complex III and Complex IV (i.e. cytochrome C oxidase) in the electron transport chain. Indeed, −Pi tobacco cells with silenced AOX undergo respiratory and growth restriction, and enhanced ROS production, relative to wild-type controls (Parsons et al., 1999). Here, we found that AOX1A (AT3G22370; +Pi: −1.16 Log_2_FC; +Phi: −0.27 Log_2_FC) protein abundance was reduced in both +Pi and +Phi cells. By contrast, protein abundance of AOX1C, a second Arabidopsis AOX isozyme, was reduced only in +Pi conditions (AT3G27620; +Pi: −1.16 Log_2_FC). Overall, we observed activation of several alternative adenylate- and/or Pi conserving pathways of cytosolic glycolysis and mitochondrial respiration in −Pi cells; however, several of these pathways seemed to be more sensitive to Pi than to Phi, suggesting a possible role in Phi-induced cellular toxicity.

### Pi and Phi induce large-scale translational and post-transcriptional regulation

In order to study the effects of Pi and Phi resupply on important biological processes, we performed an association network analysis of proteomic and phosphoproteomic data. Protein association networks exhibiting significant changes in abundance and phosphorylation status under +Pi and +Phi treatments were created based on protein-protein interaction, experimental evidence and co-expression data from the StringDB database (Figures 3, 4, and S2). The association networks for proteins changing in phosphorylation status in +Pi and +Phi samples (compared to −Pi) showed several clusters of interest. In particular, the regulation of the ribosome at the protein-level was paralleled by large increases in the phosphorylation of proteins related to ribosome biogenesis and protein synthesis (Figure 3, 4, and S2). A large proportion (40-60%) of cellular Pi in plants is allocated to nucleic acids, with the most abundant fraction of nucleic acids dedicated to the ribosome (Veneklaas et al., 2012). Indeed *Proteaceae* plants well adapted to highly Pi-impoverished soils show markedly lower levels of ribosomal RNA and ribosomal protein in developing leaves (Sulpice et al., 2014; Lambers et al., 2015). Remarkably, together with our proteomic and phosphoproteomic data, this suggests that slowing down protein synthesis through a decrease in ribosome levels and phosphorylation of translation-related proteins is a viable acclimation strategy for −Pi plants.

**Figure 3.**
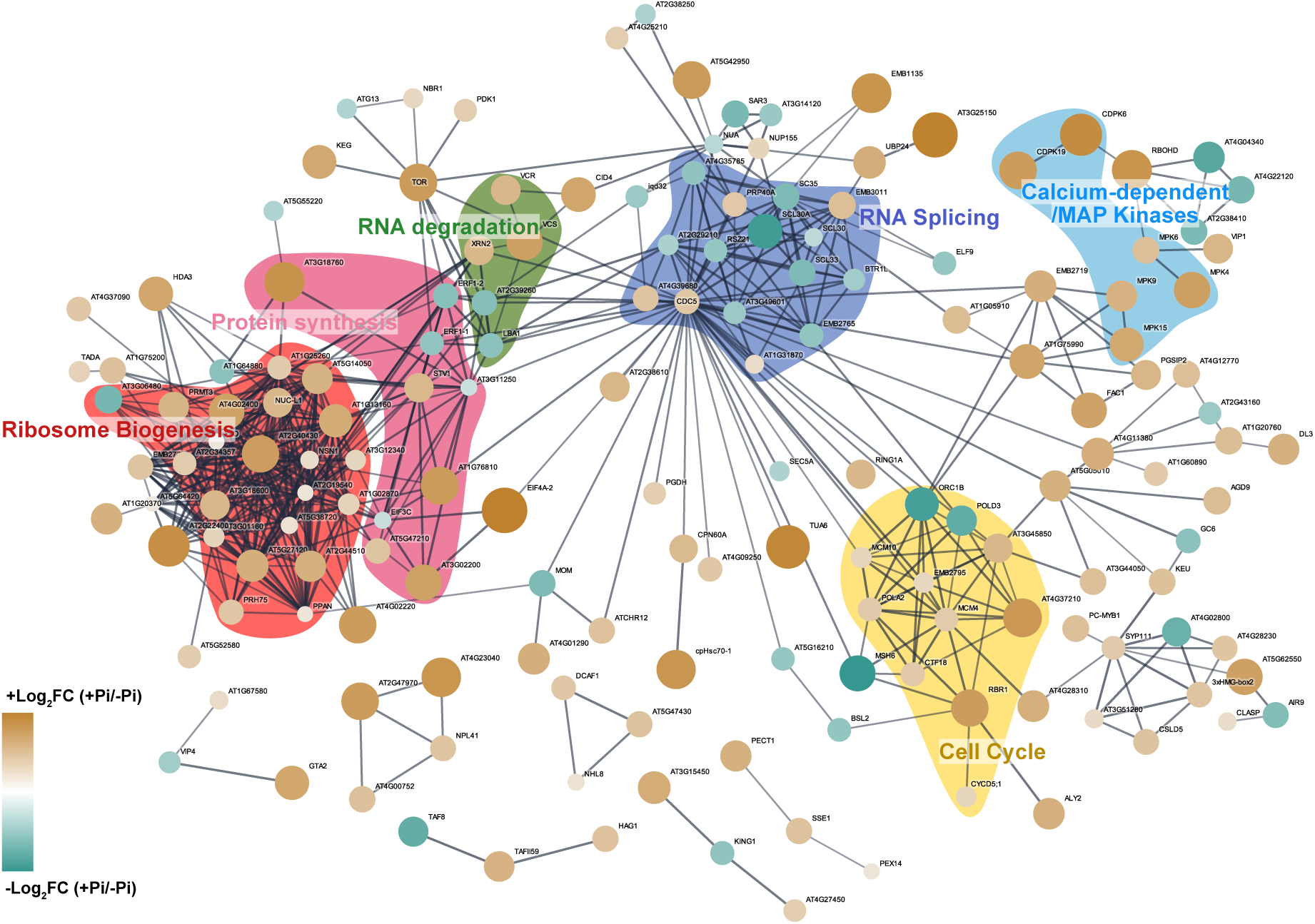
STRING-DB Association network analysis of proteins exhibiting a significant change in phosphorylation upon Pi re-feeding. Node sizes and colors are scaled by median Log_2_FC change of +Pi vs −Pi, ranging from +5.82 (Brown; large) to −4.20 (Green; small). Edge thickness represents the association confidence between two connected nodes and ranges from 0.7 to 1.0, as determined by String-DB. Highlighted regions of grouped proteins depicted the broader process relationships between associating proteins. Nodes with no connections ≥ 0.7 are not depicted.

**Figure 4.**
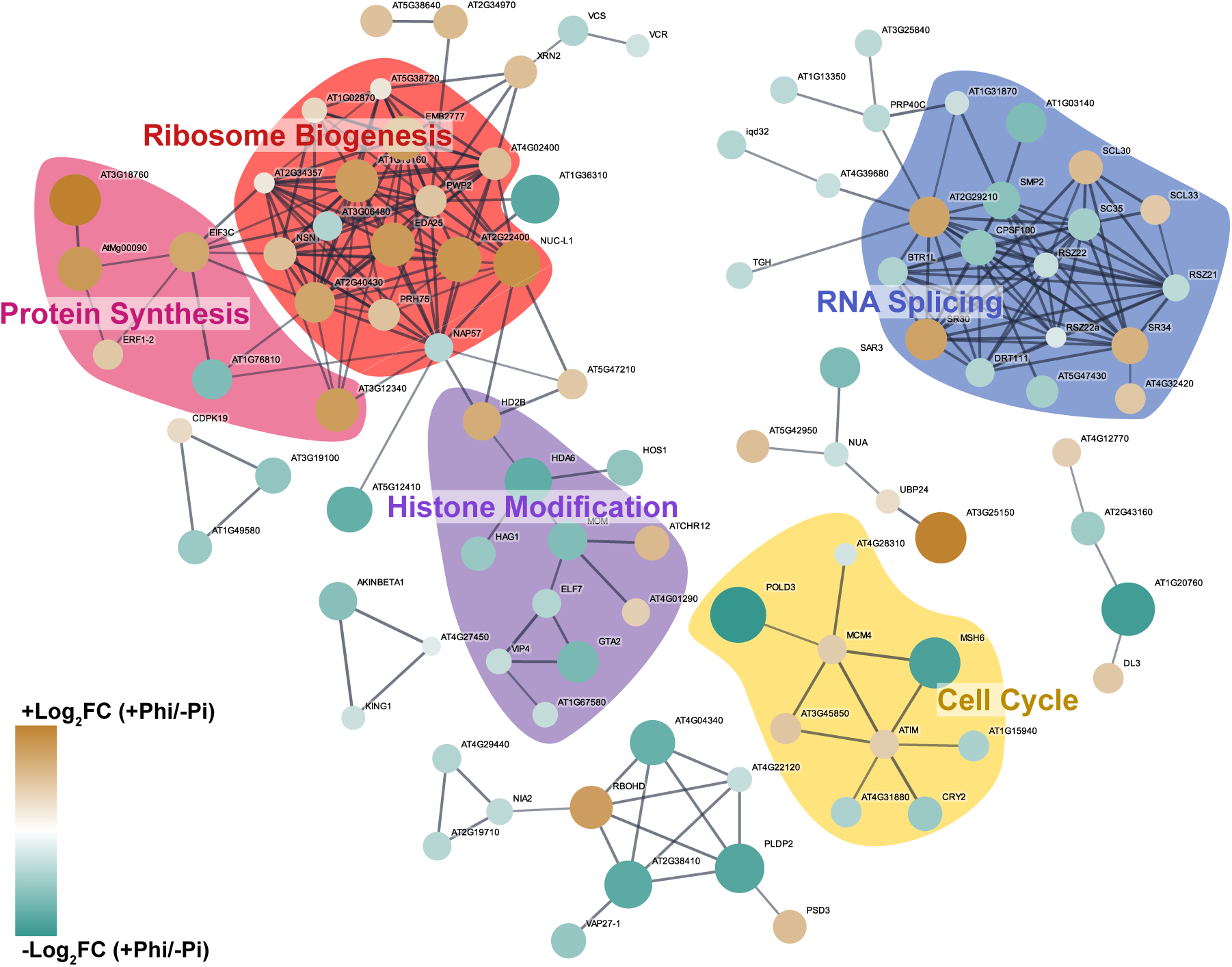
STRING-DB Association network analysis of proteins exhibiting a significant change in phosphorylation upon Phi re-feeding. Node sizes and colors are scaled by median Log_2_FC change of +Pi vs −Pi, ranging from +4.70 (Brown; large) to −5.30 (Green; small). Edge thickness represents the association confidence between two connected nodes and ranges from 0.7 to 1.0, as determined by String-DB. Highlighted regions of grouped proteins depicted the broader process relationships between associating proteins. Nodes with no connections ≥ 0.7 are not depicted.

Furthermore, our association networks of phosphoproteins changing in response to +Pi and +Phi also contain clusters of RNA splicing-related proteins. Intriguingly, and in contrast to most other processes, the phosphorylation status of these proteins was significantly reduced upon Pi- or Phi-resupply (Figures 2B, Table S2); or put another way, was increased upon Pi-starvation. Other phosphoprotein clusters of interest include calcium dependent / mitogen activated protein kinases, cell cycle proteins, and proteins involved in histone modifications and RNA degradation processes. Together, the phosphorylation-level changes in proteins involved in translation and post-transcriptional regulation suggest that Pi-starvation results in a decline in protein synthesis via combined disruption of ribosome biogenesis, translation, and RNA splicing.

### Calcium-dependent and mitogen-activated protein kinases are Pi responsive

Examination of our dataset found 70 total protein kinases that show significant changes in their phosphorylation status under changing Pi / Phi conditions, including: calcium dependent protein kinases (CDPK), mitogen activated protein kinases (MPK), receptor-like cytoplasmic kinases (RCLK) and leucine-rich repeat receptor-like kinases (LRR-RLK; Table S2). To further our understanding of how these kinases may intersect with our phosphoproteome data, we performed a phosphorylation site enrichment analysis (Table 3). An enrichment of an MPK and CDPK motif was apparent in both the +Pi and +Phi datasets, in addition to a casein kinase II- (CKII) motif also enriched in the +Pi dataset. Other currently unknown motifs were also enriched; however, their connection to specific protein kinase subfamiles requires further targeted characterization.

**Table 3.**
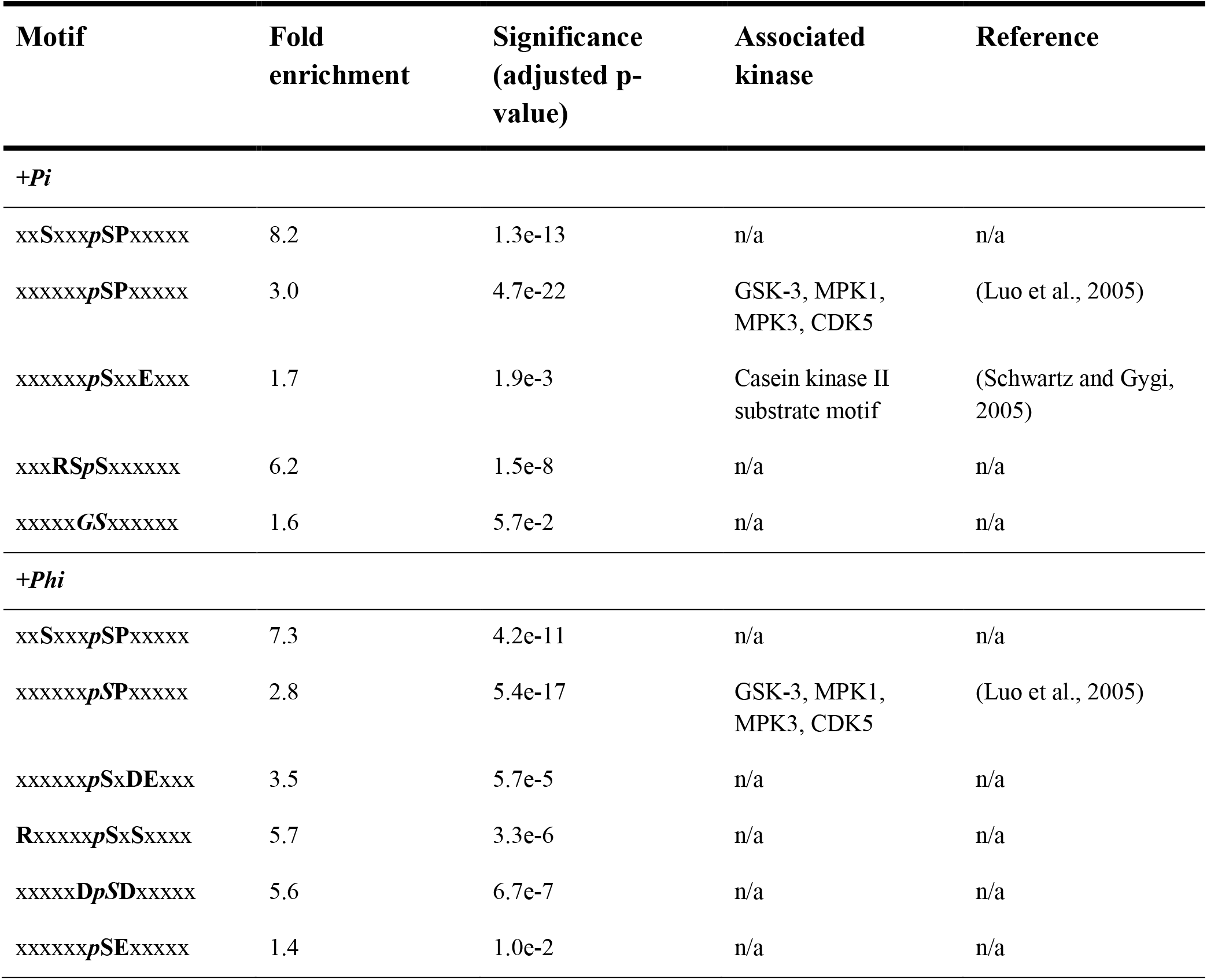
Enriched phospho-motifs identified in the significantly changing phosphoproteome of +Pi and +Phi samples compared to −Pi.

Amongst the protein kinases with annotated network associations to other proteins, we observed a number of CDPKs and MPKs with large declines in their phosphorylation status in response to +Pi (Figures 2 and 3). These include CDPK3 (AT2G17290 / CPK6; +2.82 median Log_2_FC), CDPK6 (AT4G23650 / CPK3; +4.62 median Log_2_FC), CDPK19 (AT5G19450 / CPK8, +4.49 median Log_2_FC), MPK4 (AT4G01370; +4.27 median Log_2_FC), MPK6 (AT2G43790; +2.64 median Log_2_FC), MPK9 (AT3G18040; +3.23 median Log_2_FC) and MPK15 (AT1G73670; +3.88 median Log_2_FC) (Figure 2, Table S2). Only CDPK19 (+1.24 median Log_2_FC) increased in phosphorylation in response to +Pi. Two CDPK related kinases (CDPKRK) decreased in their phosphorylation status (AT1G49580; −2.42 median Log_2_FC and AT3G19100; −2.67 median Log_2_FC) in response to +Phi (Figure 3, Table S2); while no MPKs exhibited an increase in phosphorylation under −Pi conditions. Specific examination of the significantly changing phosphorylation sites of CDPK3, 6 and 19 found that these sites fall outside the conserved canonical protein kinase activation motif (Supplemental Data 1). However, as auto-phosphorylation of the activation loop is not required for CDPK activity(Bender et al., 2018), CDPKs identified in our data may in fact be activated in response to +Pi and +Phi. Further, the observed phosphorylation events may serve to affect CDPK activity directly upon Ca^2+^ activation or by facilitating protein-protein interactions. On the other hand, each significantly fluctuating MPK phosphorylation site was found within the activation motif on amino acids with well-characterized effects (Nishimoto and Nishida, 2006), indicating that under +Pi conditions, MPK4, 6, 9 and 15 are all present in an activated state (Supplemental Data 1). Lastly, our finding that MPK and CDPKs increase their phosphorylation status in response to Pi and/or Phi refeeding (Supplemental Table 2) corroborates previous reports findings in rice (Yang et al., 2019) and maize (Li et al., 2014) that found decreases in MPK and CDPK phosphorylation during Pi starvation. When combined with our findings, this suggests that MPK and CDPK protein kinase involvement in the PSR is conserved between monocots and dicots.

CDPK3, 6 and 19 are all involved in ABA signaling (Zou et al., 2015; Li et al., 2018), which itself has a role in Pi-nutrition. Aleksza and colleagues (2017) determined that ABA can stimulate proline production during Pi-starvation through myeloblastosis (MYB) and ABA-responsive element binding protein (AREB) transcription factor mediated increases in D1-PYRROLINE-5-CARBOXYLATE SYNTHETASE 1 (P5CS1) expression levels(Aleksza et al., 2017). Here, we find ABA-RESPONSIVE ELEMENT BINDING PROTEIN 3 (AREB3; AT3G56850) phosphorylation status markedly increased (4.10 Log_2_FC) under +Pi conditions, indicating that ABA also plays a role under adequate Pi nutrition. AREB transcription factors are phosphorylated by CDPKs (Grandellis et al., 2016) and CDPK3 directly interacts with AREB3 (Zhang et al., 2020). Interestingly, multiple CDPKs were reported to associate with AREB3, suggesting that CDPK3, 6 and 19 may have specific Pi-nutrition impacts through AREB3 (Zhang et al., 2020). Furthermore, CDPKs have been implicated in the regulation of plant nitrate responses, suggesting a prominent role in nutrient signaling (Liu et al., 2017) that, as we show here, also extends to Pi nutrition. CDPK substrate motifs in plants demonstrate a high level of variability (Bender et al., 2018). Our motif enrichment analysis found a CDPK-like motif enriched under +Pi that resembles known CDPK motifs (Table 3); however, this requires further validation as a substrate motif for CDPK3, 6 and 19.

Plant MPKs have a known role in the perception and adaptation to environmental and physiological changes, including nutritional signaling (Chardin et al., 2017). Here we find that MPK members from A (MPK6), B (MPK4) and D (MPK9 and 15) subclades become activated under +Pi conditions (Figure 3; Supplemental Data 1) and their potential role under +Pi conditions supported by enrichment of a consensus MPK-like phosphorylation motif (Table 3). Of these MPK4 and MPK9 are known to be transcriptionally regulated upon −Pi conditions (Chardin et al., 2017). MPK3 and 6 are also transcriptionally induced under −Pi conditions and *mpk3* and *mpk6* plants accumulate less Pi than wild-type plants (Lei et al., 2014). Our protein-level data indicates that MPK3 and 6 are activated upon +Pi relative to −Pi conditions (Figure 3), which supports previous findings that MKK9-activated MPK3 and 6 increase Pi uptake (Lei et al 2014). MPK9 exhibits activation site specific-phosphorylation (Nagy et al., 2015) and MPK15 was reported to be active-site phosphorylated in the pollen phosphoproteome (Mayank et al., 2012).

Of the 70 protein kinases changing in our experiment, 15 are leucine rich receptor-like kinases (LRR-RLKs) (Table S2; (Zulawski et al., 2014)). Two potentially interesting candidates amongst these LRR-RLKs are PXC1 (At2g36570) and MIK2 (AT4G08850). PXC1 has roles in vascular development and cell elongation (Kim et al., 2009; Wang et al., 2013), while MIK2 has been connected to cell wall integrity sensing in response to growth and environmental cues (Van der Does et al., 2017). Overall, the multitude of largely uncharacterized LRR-RLKs and other protein kinases we have uncovered here that possess Pi-responsive phosphorylation events, each represent new opportunities to connect cell signaling through protein phosphorylation to plant cell Pi-nutrition.

### Opposing regulation in response to Pi- and Phi-resupply

A major aim of this study is to identify distinct protein-level molecular changes in response to +Pi and +Phi. In total, we only find 10 proteins that display opposing changes in abundance upon +Pi and +Phi (Figure 2A, Table S1). These changes are also relatively minor in terms of fold changes, suggesting again that specific responses to +Pi versus +Phi occur primarily at the level of protein phosphorylation. Among these proteins is GALANTHUS NIVALIS AGGLUTININ-RELATED AND APPLE DOMAIN LECTIN-1 (GAL1; AT1G78850). GAL1 expression is induced by −Pi, and the protein is dual targeted to the vacuole and cell walls of −Pi Arabidopsis suspension cells (Ghahremani et al., 2019). GAL1 also specifically interacts with and activates a high mannose glycoform of the Pi scavenging AtPAP26 (Ghahremani et al., 2019).

Overall, we observed opposing fluctuations in protein phosphorylation levels across 69 proteins (Figure 2C, Table S2). Of these, only 9 proteins showed decreased phosphorylation upon +Pi and increased phosphorylation upon +Phi. These include 5 proteins involved in RNA splicing (SCL30/ AT3G55460, SCL33/AT1G55310, AT2G29210, CFM9/AT3G27550, AT4G32420), 2 translation-related proteins (EIF3C/AT3G56150, ERF1-2/AT1G12920) and one protein kinase (AT3G58760), reinforcing the importance of phosphorylation mediated regulation of these core cellular processes as found in our association network analysis. These results suggest that the phosphorylation status of certain proteins within these large multi-subunit protein complexes may act as a molecular switch that discriminates between +Phi and +Pi (Figure 3 and 4).

Furthermore, 60 proteins in our dataset show significant changes in phosphorylation levels in opposite directions in response to +Pi and +Phi. Among these were 7 splicing related proteins, 8 protein kinases and 2 transcription factors (Table S2). Importantly, we also observed differential changes in phosphorylation of the PHT1;4 transporter (+Pi: 1.08 median Log_2_FC, +Phi: −1.56 median Log_2_FC). The differential median phosphorylation of PHT1;4 is caused by the extensive dephosphorylation of the previously characterized Ser524 site (−5.11 Log_2_FC) (Bayle et al 2011) as well as increased phosphorylation of the newly discovered Ser515 (+1.08 Log_2_FC) site upon +Pi, whereas upon +Phi, only the Ser524 site was moderately dephosphorylated (−1.56 Log_2_FC). We also observed opposing phosphorylation-level changes in HAT1 (AT3G54610), a histone acetyltransferase that regulates the transcription of a long non-coding RNA *At4* that in turn regulates the expression of miR399 and hence its target the PHO2 protein (Wang et al., 2019). PHO2 is a ubiquitin conjugating E2 enzyme that facilitates degradation of the PHO1 Pi transporter in +Pi cells and is a well-studied player in the plant PSR (Bari et al., 2006; Liu et al., 2012). A significant increase in HAT1 phosphorylation at Ser76 upon Pi refeeding was also detected (+2.28 Log_2_FC), whereas a significant decrease in phosphorylation at the same site (−2.47 Log_2_FC) occurred upon +Phi. An additional 3 sites, Thr75, Ser84, and Ser90 (+2.28, +3.20, +2.84 Log_2_FC, respectively) were also significantly more phosphorylated in only the +Pi-fed cells and not in +Phi cells. The *At4/IPS1*-miR399-PHO2/PHO1 pathway is among the best-studied PSR of Arabidopsis (Oldroyd and Leyser, 2020) and our results suggest an important upstream role for HAT1 in this pathway via +Pi- and +Phi-specific phosphorylation changes. Since we did not detect changes in any of the other protein members of the *At4/IPS1*-miR399-PHO2/PHO1 pathway such as MYB2, WRKY6, PHO2 and PHO1 at either the protein abundance or phosphorylation levels, further investigation of HAT1 post-translational regulation in this pathway is needed.

## CONCLUSIONS

Here we present an in-depth quantitative proteomic and phosphoproteomic analysis of the Arabidopsis phosphate starvation response. Our results provide exciting new global insights into protein abundance and signalling changes in plants responding to Pi and Phi (Figure 5). In particular, we validated and significantly expanded our current understanding of the PSR through the quantification of abundance changes in several canonical Pi starvation inducible proteins such as the PAPs, PHTs, PHF1, and phospholipases. Our data provides further support to earlier hypotheses in the field that phosphate availability regulates the functioning of metabolic bypasses to adenylate-dependent reactions in plants (including the phosphorylation of a key GABA-shunt enzyme) in phosphate sensing. We also provide direct evidence that responses to Pi nutrition, rather than Pi sensing or transport (as mimicked by Phi), occur primarily at the level of protein phosphorylation rather than protein abundance. This is highlighted by our elucidation of the differential phospho-regulation of PHT1;4 and HAT1 in response to Pi / Phi, which provide important additions to well-characterized aspects of the PSR such as *At4/IPS1*-miR399-PHO2/PHO1 pathway. Additionally, both Pi and Phi induced large-scale fluctuations in the phosphorylation status of translation and RNA splicing proteins, which supports the emerging idea that regulation of mRNA splicing is a generalized stress response coupled with reduced mRNA translation (Chaudhary et al., 2019; Morgan et al., 2019; Parenteau et al., 2019). For example, 12 of the 69 proteins that exhibited opposing changes in their phosphorylation status between Pi/Phi-fed and −Pi cells are splicing related proteins. RNA splicing has not yet been directly studied in the context of Pi-starvation; however, our results provide a clear indication of a role for the post-translational regulation of RNA splicing in response to Pi-starvation. We also identify Pi-associated phosphorylation changes in a number of MAP-kinases and CDPKs, generating important targets for future work to elucidate intracellular signalling pathways responsive to Pi nutrition. Given the relative paucity of research to date into protein phosphorylation during the PSR, our findings call for a greater focus on Pi-responsive kinase signalling in future studies. Overall, we provide extensive protein-level insights into the Pi starvation and Pi / Phi re-feeding responses that not only increases our fundamental understanding of plant nutrition-stress responses but will also assist in the development of crops with increased phosphate acquisition and use efficiency.

**Figure 5.**
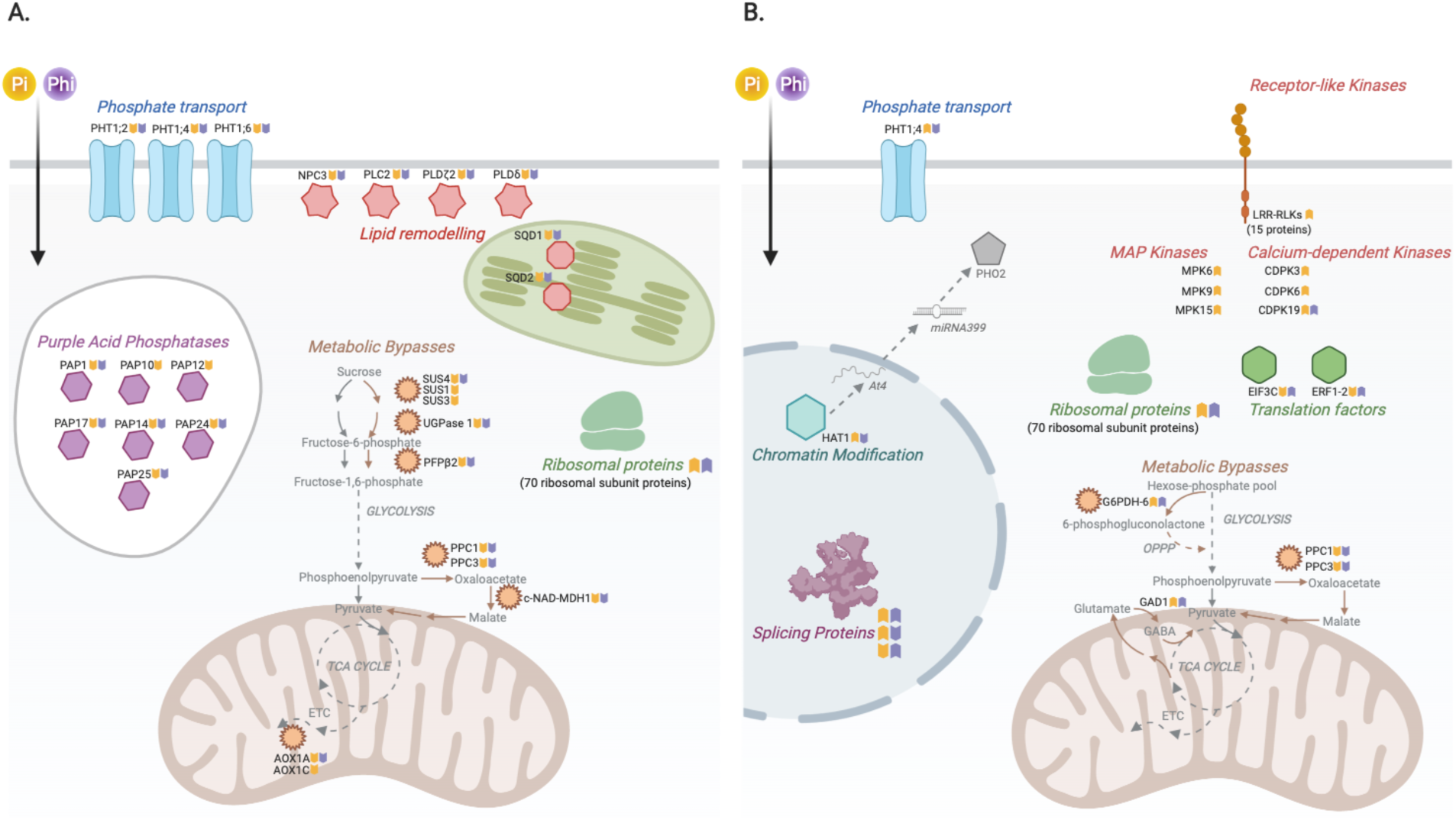
Summary of observed changes in (A) protein abundance and (B) protein phosphorylation upon Pi and Phi re-feeding. PHT: PHOSPHATE TRANSPORTER; PAP: PURPLE ACID PHOSPHATASE; SQD: SULFOQUINOVOSYLDIACYLGLYCEROL; SUS: SUCROSE SYNTHASE; UGPase:UDP-GLUCOSE PYROPHOSPHORYLASE; PFPβ2: PHOSPHOFRUCTOKINASE BETA 2; PPC: PHOSPHOENOLPYRUVATE CARBOXYLASE; c-NAD-MDH1: CYTOSOLIC NAD-MALATE DEHYDROGENASE; AOX: ALTERNATIVE OXIDASE; LRR-RLK: LEUCINE-RICH REPEAT RECEPTOR-LIKE KINASE; MPK: MITOGEN ASSOCIATED KINASE; CDPK: CALCIUM DEPENDENT PROTEIN KINASE; HAT1: HISTONE ACETYLTRANSFERASE 1; PHO2: PHOSPHATE 2; EIF3C: ELONGATION INITIATION FACTOR 3C; ERF1-2: ELONGATION RELEASE FACTOR 1-2; G6PDH: GLUCOSE-6-PHOSPHATE DEHYDROGENASE; GAD1: GLUTAMATE DECARBOXYLASE 1; TCA: Tricarboxylic Acid; ETC: Electron Transport Chain.

## MATERIALS AND METHODS

### Plant material

Heterotrophic Arabidopsis (*Arabidopsis thaliana*, cv. Landsberg erecta) suspension cells were maintained in standard Murashige-Skoog media at 21 °C in the dark as previously described (Veljanovski et al., 2006). For the generation of experimental samples, 30 mL aliquots of cell suspension that had been subcultured for 6 days in media containing 5 mM K_2_HPO_4_ were used to inoculate 12 separate 500 mL flasks that each contained 120 mL of fresh media lacking Pi. Three days later, the cells were fed sterilized 0 or 2 mM Pi, or 2 mM Phi (*n* = 4 each). Cells were harvested 2 days later by vacuum filtration and stored at –80 °C.

### Preparation of clarified extracts

Quick-frozen cells were ground to a powder in liquid N_2_ and homogenized (1:2, w/v) using a mortar and pestle and a small scoop of sand in ice-cold Buffer A which contained 50 mM imidazole (pH 7.0), 1 mM EDTA, 2 mM MgCl_2_, 25 mM NaF, 1 mM NaVO_3_^−^, 1 mM NaMoO_4_, 50 nM microcystin-LR, 0.1% (v/v) Triton X-100, 10% (v/v) glycerol, 1% (w/v) polyvinyl(polypyrrolidone), 1 mM phenylmethylsulfonyl fluoride, and 2.5 μL/ml ProteCEASE-100 (G-Biosciences). Extracted proteins were filtered through Miracloth before centrifuging at 14,000 xg for 10 min at 4 °C. The resulting clarified extracts were rapidly prepared for SDS/PAGE and immunoblotting or assayed for total protein and APase activity.

### Acid phosphatase assay and protein concentration determination

APase activity was measured by coupling the hydrolysis of phosphoenolpyruvate to pyruvate to the lactate dehydrogenase reaction and assaying at 24 °C by monitoring the oxidation of NADH at 340 nm using a Molecular Devices Spectromax Plus Microplate spectrophotometer. Optimal APase assay conditions were: 50 mM Na-acetate (pH 5.6), 5 mM phosphoenolpyruvate, 0.2 mM NADH, and 3 units/ml of rabbit muscle lactate dehydrogenase in a final volume of 0.2 ml. Assays were corrected for any background NADH oxidation by omitting phosphoenolpyruvate from the reaction mixture. All APase assays were linear with respect to time and concentration of enzyme assayed. One unit of activity is defined as the amount resulting in the utilization of 1 μmol/min of substrate. Protein concentrations were determined using the Coomassie Blue G-250 dye binding method with bovine γ-globulin as the protein standard.

### Immunoblotting

SDS/PAGE was performed using Bio-Rad Protean III mini-gel (200 V, 50 min) as previously described (Gregory et al., 2009). Immunoblotting was conducted by electroblotting proteins from gels onto PVDF membranes, and immunoreactive polypeptides were visualized using an alkaline-phosphatase conjugated secondary antibody with chromogenic detection. Immunoblots were probed with anti-(*Ricinus communis* PEPC) immune serum (anti-PEPC) that had been raised in rabbits against native class-1 PEPC (RcPPC3) fully purified from developing castor beans (Gennidakis et al., 2007). Anti-(phosphorylation site-specific)-IgG (anti-pSer11) immunoblots were probed with a polyclonal antibody raised against a synthetic phosphopeptide matching RcPPC3’s conserved N-terminal Ser11 phosphorylation domain (L-E-K-L-A-pS-I-D-A-Q-L-R) (Tripodi et al., 2005). For anti-pSer11 immunoblots the corresponding de-phosphopeptide (10 μg/mL) was used to block any nonspecific cross reaction with non-phosphorylated PEPC polypeptides (Tripodi et al., 2005). All immunoblot results were replicated at least two times, with representative results shown in the various figures.

### Mass spectrometry sample preparation and phosphopeptide enrichment

Quick-frozen cells were lyophilized, then ground to a fine powder under liquid N_2_ and extracted using filter-assisted sample preparation with a 30K MWCO Amicon filter units as described by Uhrig et al. (2019). Digested peptides (500 μg) were desalted using C18 Sep-Pak Cartridges (Waters), followed by immediate enrichment of phosphorylated peptides using NP-Ti0_2_ (Sachtopore; SNX 030S 005 #9205/1639) as previously described (Wiśniewski et al., 2009; Uhrig et al., 2019). Unbound peptides and NP-Ti02 enriched Ti02 peptides were dried, dissolved in 3% ACN/0.1% TFA, desalted using ZipTip C18 pipette tips (ZTC18S960; Millipore), then dried and dissolved in 3.0% ACN/0.1% FA prior to MS analysis.

### LC-MS/MS analysis

Peptide and phosphopeptide samples were analyzed using a QExactive Orbitrap mass spectrometer (Thermo Scientific). Dissolved samples were injected using an Easy-nLC 1000 system (Thermo Scientific) and separated on 50 cm ES803 Easy-Spray PepMap C18 Column (Thermo Scientific). The column was equilibrated with 100% solvent A (0.1% formic acid (FA) in water). Peptides were eluted using the following gradient of solvent B (0.1% FA in ACN): 5% to 22% B, 0 – 110 min; 22% - 35% B, 110 – 120 min; 35 - 95% B, 120 – 125 min at a flow rate of 0.3 μl/min at 50°C. High accuracy mass spectra were acquired in data-dependent acquisition mode. All precursor signals were recorded in a mass range of 300 - 1700 m/z and a resolution of 70000 at 200 m/z. The maximum accumulation time for a target value of 3e6 was set to 120 ms. Up to 12 data dependent MS/MS were recorded using quadrupole isolation with a window of 2 Da and HCD fragmentation with 30% fragmentation energy. A target value of 1e6 was set for MS/MS using a maximum injection time of 250 ms and a resolution of 70000 at 200 m/z. Precursor signals were selected for fragmentation with charge states from +2 to +7 and a signal intensity of at least 1e5. All precursor signals selected for MS/MS were dynamically excluded for 30 s.

### Label-free quantitation

Raw data were processed using MaxQuant software version 1.6.14 (https://www.maxquant.org/; (Cox and Mann, 2008; Tyanova et al., 2016)). MS/MS spectra were searched with the Andromeda search engine against a decoyed (reversed) version of the Arabidopsis protein database from TAIR (release TAIR10) concatenated with a collection of 261 known mass spectrometry contaminants. Trypsin specificity was set to one missed cleavage and a protein and PSM false discovery rate of 1% was deployed. Minimal peptide length was set to seven and Match between runs along with requantify options enabled. Fixed modifications included: Carbamidomethylation of cysteine residues, while variable modifications included: methionine oxidation (both proteome and phosphoproteome analyses) and phosphorylated serine, threonine and tyrosine (pSTY) for analysis of the phosphoproteome.

### Bioinformatic analysis

Downstream data analyses were performed using Perseus version 1.6.10.43 (Tyanova et al., 2016). For both proteome and phosphoproteome analysis, reverse hits and contaminants were removed, the data Log_2_-transformed, followed by a data subselection criteria of n=2 of 4 replicates in at least one sample. Missing values were replaced using the normal distribution imputation method. For the phosphoproteome, phosphorylation sites were filtered for a localization probability of ≥ 0.75 prior to quantification. Subsequently, significantly changing differentially abundant proteins and phosphorylation sites were determined and corrected for multiple comparisons (Benjamini-Hochberg FDR *p-value* < *0.05; q-value*). Overall fold changes in phosphorylation status of individual phosphoproteins were calculated as the median Log_2_ fold changes of every unique phosphopeptide in the respective protein.

Association networks presented in Figures 3, 4 and S2 were constructed in Cytoscape version 3.7.2 using StringDB databases (co-expression, experimental and interaction data) with a minimum confidence threshold of 0.7. The size of each node was scaled with absolute Log2 fold change values and a color scale corresponding to Log_2_ fold change values was applied to each node. Clusters were manually defined and annotated based on inspection of node protein descriptions and based on gene ontology.

For identification of enriched phosphosite motifs, we created a custom program called PSMFinder to isolate peptide windows and subsequently perform motif enrichment analysis. PSMFinder first isolates 14AA windows centered on the modified amino acids (STY) in our lists of significantly changing phosphopeptides (Table S2). The isolated 14AA peptides for +Pi and +Phi datasets were next analysed using a standalone installation of the MoMo program (Cheng et al., 2019) with the motif-x algorithm (Schwartz and Gygi, 2005) in Meme Suite 5.1.1 (Bailey et al., 2009) using the following settings:- width: 13, remove unknowns: false, minimum number of motif occurrences: 10 and score threshold of 0.0001.

### Data availability

Raw data have been deposited to the ProteomeExchange Consortium (https://proteomecentral.proteomexchange.org) via the PRIDE partner repository with the dataset identifiers PXD018989. Custom scripts used for data analysis and plotting, as well as program outputs are provided under a GNU Affero General Public License v3.0 license at https://github.com/UhrigLab/mehta2020-Pi-proteome.

## ACKNOWLEDGEMENTS

The authors would like to thank Dr. Bernd Roschitzki of the Functional Genomics Center Zurich for useful discussions. This project was supported by National Science and Engineering Research Council of Canada (NSERC) research and equipment grants to RGU and WCP, a Queen’s Research Chair award to WCP, and an OECD CRP Travel Fellowship to MP-F. DM and MG were funded by a Swiss National Science Foundation Early Postdoc Mobility grant (181602) and an Ontario Trillium Scholarship, respectively.

## SUPPORTING INFORMATION

**Supporting Data 1. Significantly changing phosphorylation sites mapped onto the corresponding CDPK and MAPK proteins.** Full-length protein sequences were obtained from Phytozome v12 (https://phytozome.jgi.doe.gov). Highlighted in yellow is the significantly changing phosphorylated peptide(s) identified, while amino acids bold and in red represent the corresponding phosphorylation sites (site ID ≥ 0.75).

**Table S1. Lists of proteins that were quantified and have significant changes in abundance across +Pi, +Phi and −Pi.**

**Table S2. Lists of proteins that were quantified and have significant changes in phosphorylation across +Pi, +Phi and −Pi.**

**Figure S1. Subcellular localization of proteins exhibiting a significant change in abundance or phosphorylation status in response to Pi or Phi.** All significantly changing proteins were subjected to consensus subcellular localization analysis using SUBA4 (https://suba.live/;. Data were plotted as a percent of the total significantly changing species in response to re-feeding with either Pi or Phi.

**Figure S2. Association network analysis of Pi induced changes in the Arabidopsis proteome.** (A) Node sizes and colors are scaled by median Log_2_FC change of +Pi vs −Pi, ranging from +4.20 (Brown; large) to −7.13 (Green; small) Edge thickness represents the association confidence between two connected nodes and ranges from 0.7 to 1.0, as determined by String-DB. (B) Node sizes and colors are scaled by median Log_2_FC change of +Phi vs −Pi, ranging from +3.20 (Brown; large) to −1.5 (Green; small). Edge thickness represents the association confidence between two connected nodes and ranges from 0.7 to 1.0, as determined by String-DB. Highlighted regions of grouped proteins depicted the broader process relationships between associating proteins. Nodes with no connections ≥ 0.7 are not depicted.

## References

Aleksza D, Horváth GV, Sándor G, Szabados L (2017) Proline Accumulation Is Regulated by Transcription Factors Associated with Phosphate Starvation. Plant Physiol 175: 555–567

Bailey TL, Boden M, Buske FA, Frith M, Grant CE, Clementi L, Ren J, Li WW,Noble WS(2009) MEME SUITE: tools for motif discovery and searching. Nucleic Acids Res 37: W202–8

Bari R, Datt Pant B, Stitt M, Scheible W-R (2006) PHO2, microRNA399, and PHR1 define a phosphate-signaling pathway in plants. Plant Physiol 141: 988–999

Bown AW, Shelp BJ (2020) Does the GABA shunt regulate cytosolic GABA? Trends Plant Sci 25: 422–424

Carswell C, Grant BR, Theodorou ME, Harris J, Niere JO, Plaxton WC (1996) The Fungicide Phosphonate Disrupts the Phosphate-Starvation Response in Brassica nigra Seedlings. Plant Physiol 110: 105–110

Carswell MC, Grant BR, Plaxton WC (1997) Disruption of the phosphate-starvation response of oilseed rape suspension cells by the fungicide phosphonate. Planta 203: 67–74

Chardin C, Schenk ST, Hirt H, Colcombet J, Krapp A (2017) Review: Mitogen-Activated Protein Kinases in nutritional signaling in Arabidopsis. Plant Sci 260: 101–108

Chaudhary S, Jabre I, Reddy ASN, Staiger D, Syed NH (2019) Perspective on alternative splicing and proteome complexity in plants. Trends Plant Sci 24: 496–506

Cheng A, Grant CE, Noble WS, Bailey TL (2019) MoMo: discovery of statistically significant post-translational modification motifs. Bioinformatics 35: 2774–2782

Cordell D, White S (2014) Life’s bottleneck: sustaining the world’s phosphorus for a food secure future. Annu Rev Environ Resour 39: 161–188

Cox J, Mann M (2008) MaxQuant enables high peptide identification rates, individualized p.p.b.-range mass accuracies and proteome-wide protein quantification. Nat Biotechnol 26: 1367–1372

Del Vecchio HA, Ying S, Park J, Knowles VL, Kanno S, Tanoi K, She Y-M, Plaxton WC (2014) The cell wall-targeted purple acid phosphatase AtPAP25 is critical for acclimation of Arabidopsis thaliana to nutritional phosphorus deprivation. Plant J 80: 569–581

Duan G, Walther D, Schulze WX (2013) Reconstruction and analysis of nutrient-induced phosphorylation networks in Arabidopsis thaliana. Front Plant Sci 4: 540

Engelsberger WR, Schulze WX (2012) Nitrate and ammonium lead to distinct global dynamic phosphorylation patterns when resupplied to nitrogen-starved Arabidopsis seedlings. Plant J 69: 978–995

Essigmann B, Güler S, Narang RA, Linke D, Benning C (1998) Phosphate availability affects the thylakoid lipid composition and the expression of SQD1, a gene required for sulfolipid biosynthesis in Arabidopsis thaliana. Proc Natl Acad Sci USA 95: 1950–1955

Gaude N, Nakamura Y, Scheible W-R, Ohta H, Dörmann P (2008) Phospholipase C5 (NPC5) is involved in galactolipid accumulation during phosphate limitation in leaves of Arabidopsis. Plant J 56: 28–39

Gennidakis S, Rao S, Greenham K, Uhrig RG, O’Leary B, Snedden WA, Lu C, Plaxton WC (2007) Bacterial- and plant-type phosphoenolpyruvate carboxylase polypeptides interact in the hetero-oligomeric Class-2 PEPC complex of developing castor oil seeds. Plant J 52: 839–849

Ghahremani M, Tran H, Biglou SG, O’Gallagher B, She Y-M, Plaxton WC (2019) A glycoform of the secreted purple acid phosphatase AtPAP26 co-purifies with a mannose-binding lectin (AtGAL1) upregulated by phosphate-starved Arabidopsis. Plant Cell Environ 42: 1139–1157

González E, Solano R, Rubio V, Leyva A, Paz-Ares J (2005) PHOSPHATE TRANSPORTER TRAFFIC FACILITATOR1 is a plant-specific SEC12-related protein that enables the endoplasmic reticulum exit of a high-affinity phosphate transporter in Arabidopsis. Plant Cell 17: 3500–3512

Grandellis C, Fantino E, Muñiz García MN, Bialer MG, Santin F, Capiati DA, Ulloa RM (2016) Stcdpk3 phosphorylates in vitro two transcription factors involved in GA and ABA signaling in potato: strsg1 and stabf1. PLoS One 11: e0167389

Gregory AL, Hurley BA, Tran HT, Valentine AJ, She Y-M, Knowles VL, Plaxton WC (2009) In vivo regulatory phosphorylation of the phosphoenolpyruvate carboxylase AtPPC1 in phosphate-starved Arabidopsis thaliana. Biochem J 420: 57–65

Hooper CM, Castleden IR, Tanz SK, Aryamanesh N, Millar AH (2017) SUBA4: the interactive data analysis centre for Arabidopsis subcellular protein locations. Nucleic Acids Res 45: D1064–D1074

Jin J, Pawson T (2012) Modular evolution of phosphorylation-based signalling systems. Philos Trans R Soc Lond B, Biol Sci 367: 2540–2555

Jost R, Pharmawati M, Lapis-Gaza HR, Rossig C, Berkowitz O, Lambers H, Finnegan PM (2015) Differentiating phosphate-dependent and phosphate-independent systemic phosphate-starvation response networks in Arabidopsis thaliana through the application of phosphite. J Exp Bot 66: 2501–2514

Karthikeyan AS, Varadarajan DK, Mukatira UT, D’Urzo MP, Damsz B, Raghothama KG (2002) Regulated expression of Arabidopsis phosphate transporters. Plant Physiol 130: 221–233

Kim S, Kim S-J, Shin Y-J, Kang J-H, Kim M-R, Nam KH, Lee M-S, Lee S-H, Kim Y-H, Hong S-K, et al (2009) An atypical soybean leucine-rich repeat receptor-like kinase, GmLRK1, may be involved in the regulation of cell elongation. Planta 229: 811–821

Kuo H-F, Hsu Y-Y, Lin W-C, Chen K-Y, Munnik T, Brearley CA, Chiou T-J (2018) Arabidopsis inositol phosphate kinases IPK1 and ITPK1 constitute a metabolic pathway in maintaining phosphate homeostasis. Plant J. doi: 10.1111/tpj.13974

Lambers H, Finnegan PM, Jost R, Plaxton WC, Shane MW, Stitt M (2015) Phosphorus nutrition in Proteaceae and beyond. Nat Plants 1: 15109

Lambers H, Plaxton WC (2015) Phosphorus: back to the roots. In WC Plaxton, H Lambers, eds, Annual plant reviews volume 48: phosphorus metabolism in plants. John Wiley & Sons, Inc., Hoboken, NJ, USA, pp 1–22

Lan P, Li W, Schmidt W (2012) Complementary proteome and transcriptome profiling in phosphate-deficient Arabidopsis roots reveals multiple levels of gene regulation. Mol Cell Proteomics 11: 1156–1166

Lei L, Li Y, Wang Q, Xu J, Chen Y, Yang H, Ren D (2014) Activation of MKK9-MPK3/MPK6 enhances phosphate acquisition in Arabidopsis thaliana. New Phytol 203: 1146–1160

Li K, Xu C, Fan W, Zhang H, Hou J, Yang A, Zhang K (2014) Phosphoproteome and proteome analyses reveal low-phosphate mediated plasticity of root developmental and metabolic regulation in maize (Zea mays L.). Plant Physiol Biochem 83: 232–242

Li K, Xu C, Li Z, Zhang K, Yang A, Zhang J (2008) Comparative proteome analyses of phosphorus responses in maize (Zea mays L.) roots of wild-type and a low-P-tolerant mutant reveal root characteristics associated with phosphorus efficiency. Plant J 55: 927–939

Li Z, Takahashi Y, Scavo A, Brandt B, Nguyen D, Rieu P, Schroeder JI (2018) Abscisic acid-induced degradation of Arabidopsis guanine nucleotide exchange factor requires calcium-dependent protein kinases. Proc Natl Acad Sci USA 115: E4522–E4531

Liang C-Y, Chen Z-J, Yao Z-F, Tian J, Liao H (2012) Characterization of two putative protein phosphatase genes and their involvement in phosphorus efficiency in Phaseolus vulgaris. J Integr Plant Biol 54: 400–411

Lim H, Cho M-H, Jeon J-S, Bhoo SH, Kwon Y-K, Hahn T-R (2009) Altered expression of pyrophosphate: fructose-6-phosphate 1-phosphotransferase affects the growth of transgenic Arabidopsis plants. Mol Cells 27: 641–649

Liu K-H, Niu Y, Konishi M, Wu Y, Du H, Sun Chung H, Li L, Boudsocq M, McCormack M, Maekawa S, et al (2017) Discovery of nitrate-CPK-NLP signalling in central nutrient-growth networks. Nature 545: 311–316

Liu T-Y, Huang T-K, Tseng C-Y, Lai Y-S, Lin S-I, Lin W-Y, Chen J-W, Chiou T-J (2012) PHO2-dependent degradation of PHO1 modulates phosphate homeostasis in Arabidopsis. Plant Cell 24: 2168–2183

López-Arredondo DL, Herrera-Estrella L (2012) Engineering phosphorus metabolism in plants to produce a dual fertilization and weed control system. Nat Biotechnol 30: 889–893

Luo M, Reyna S, Wang L, Yi Z, Carroll C, Dong LQ, Langlais P, Weintraub ST, Mandarino LJ (2005) Identification of insulin receptor substrate 1 serine/threonine phosphorylation sites using mass spectrometry analysis: regulatory role of serine 1223. Endocrinology 146: 4410–4416

Masakapalli SK, Bryant FM, Kruger NJ, Ratcliffe RG (2014) The metabolic flux phenotype of heterotrophic Arabidopsis cells reveals a flexible balance between the cytosolic and plastidic contributions to carbohydrate oxidation in response to phosphate limitation. Plant J 78: 964–977

Matthus E, Wilkins KA, Swarbreck SM, Doddrell NH, Doccula FG, Costa A, Davies JM (2019) Phosphate Starvation Alters Abiotic-Stress-Induced Cytosolic Free Calcium Increases in Roots. Plant Physiol 179: 1754–1767

Mayank P, Grossman J, Wuest S, Boisson-Dernier A, Roschitzki B, Nanni P, Nühse T, Grossniklaus U (2012) Characterization of the phosphoproteome of mature Arabidopsis pollen. Plant J 72: 89–101

McDonald AE, Grant BR, Plaxton WC (2001) Phosphite (phosphorous acid): its relevance in the environment and agriculture and influence on plant phosphate starvation response. J Plant Nutr 24: 1505–1519

Menz J, Li Z, Schulze WX, Ludewig U (2016) Early nitrogen-deprivation responses in Arabidopsis roots reveal distinct differences on transcriptome and (phospho-) proteome levels between nitrate and ammonium nutrition. Plant J 88: 717–734

Mergner J, Frejno M, List M, Papacek M, Chen X, Chaudhary A, Samaras P, Richter S, Shikata H, Messerer M, et al (2020) Mass-spectrometry-based draft of the Arabidopsis proteome. Nature 579: 409–414

Michaeli S, Fromm H (2015) Closing the loop on the GABA shunt in plants: are GABA metabolism and signaling entwined? Front Plant Sci 6: 419

Morgan JT, Fink GR, Bartel DP (2019) Excised linear introns regulate growth in yeast. Nature 565: 606–611

Nagy SK, Darula Z, Kállai BM, Bögre L, Bánhegyi G, Medzihradszky KF, Horváth GV, Mészáros T (2015) Activation of AtMPK9 through autophosphorylation that makes it independent of the canonical MAPK cascades. Biochem J 467: 167–175

Nakamura Y, Awai K, Masuda T, Yoshioka Y, Takamiya K, Ohta H (2005) A novel phosphatidylcholine-hydrolyzing phospholipase C induced by phosphate starvation in Arabidopsis. J Biol Chem 280: 7469–7476

Namchuk M, Lindsay L, Turck CW, Kanaani J, Baekkeskov S (1997) Phosphorylation of serine residues 3, 6, 10, and 13 distinguishes membrane anchored from soluble glutamic acid decarboxylase 65 and is restricted to glutamic acid decarboxylase 65alpha. J Biol Chem 272: 1548–1557

Nasr Esfahani M, Kusano M, Nguyen KH, Watanabe Y, Ha CV, Saito K, Sulieman S, Herrera-Estrella L, Tran LS (2016) Adaptation of the symbiotic Mesorhizobium-chickpea relationship to phosphate deficiency relies on reprogramming of whole-plant metabolism. Proc Natl Acad Sci USA 113: E4610–9

Nishimoto S, Nishida E (2006) MAPK signalling: ERK5 versus ERK1/2. EMBO Rep 7: 782–786

Okazaki Y, Otsuki H, Narisawa T, Kobayashi M, Sawai S, Kamide Y, Kusano M, Aoki T, Hirai MY, Saito K (2013) A new class of plant lipid is essential for protection against phosphorus depletion. Nat Commun 4: 1510

Oldroyd GED, Leyser O (2020) A plant’s diet, surviving in a variable nutrient environment. Science. doi: 10.1126/science.aba0196

O’Leary B, Fedosejevs ET, Hill AT, Bettridge J, Park J, Rao SK, Leach CA, Plaxton WC(2011a) Tissue-specific expression and post-translational modifications of plant- and bacterial-type phosphoenolpyruvate carboxylase isozymes of the castor oil plant, Ricinus communis L. J Exp Bot 62: 5485–5495

O’Leary B, Park J, Plaxton WC(2011b) The remarkable diversity of plant PEPC (phosphoenolpyruvate carboxylase): recent insights into the physiological functions and post-translational controls of non-photosynthetic PEPCs. Biochem J 436: 15–34

Pan W, Wu Y, Xie Q (2019) Regulation of ubiquitination is central to the phosphate starvation response. Trends Plant Sci 24: 755–769

Parenteau J, Maignon L, Berthoumieux M, Catala M, Gagnon V, Abou Elela S (2019) Introns are mediators of cell response to starvation. Nature 565: 612–617

Parsons HL, Yip JY, Vanlerberghe GC (1999) Increased respiratory restriction during phosphate-limited growth in transgenic tobacco cells lacking alternative oxidase. Plant Physiol 121: 1309–1320

Plaxton WC, Shane MW (2015) The Role of Post-Translational Enzyme Modifications in the Metabolic Adaptations of Phosphorus-Deprived Plants. In WC Plaxton, H Lambers, eds, Annual plant reviews volume 48: phosphorus metabolism in plants. John Wiley & Sons, Inc., Hoboken, NJ, USA, pp 99–123

Plaxton WC, Tran HT (2011) Metabolic adaptations of phosphate-starved plants. Plant Physiol 156: 1006–1015

Rouached H, Arpat AB, Poirier Y (2010) Regulation of phosphate starvation responses in plants: signaling players and cross-talks. Mol Plant 3: 288–299

Saito S, Uozumi N (2020) Calcium-Regulated Phosphorylation Systems Controlling Uptake and Balance of Plant Nutrients. Front Plant Sci 11: 44

Schwartz D, Gygi SP (2005) An iterative statistical approach to the identification of protein phosphorylation motifs from large-scale data sets. Nat Biotechnol 23: 1391–1398

Shane MW, Fedosejevs ET, Plaxton WC (2013) Reciprocal control of anaplerotic phosphoenolpyruvate carboxylase by in vivo monoubiquitination and phosphorylation in developing proteoid roots of phosphate-deficient harsh hakea. Plant Physiol 161: 1634–1644

Sharma K, D’Souza RCJ, Tyanova S, Schaab C, Wiśniewski JR, Cox J, Mann M (2014) Ultradeep human phosphoproteome reveals a distinct regulatory nature of Tyr and Ser/Thr-based signaling. Cell Rep 8: 1583–1594

Singh VK, Wood SM, Knowles VL, Plaxton WC (2003) Phosphite accelerates programmed cell death in phosphate-starved oilseed rape (Brassica napus) suspension cell cultures. Planta 218: 233–239

Su Y, Li M, Guo L, Wang X (2018) Different effects of phospholipase Dζ2 and non-specific phospholipase C4 on lipid remodeling and root hair growth in Arabidopsis response to phosphate deficiency. Plant J 94: 315–326

Sulpice R, Ishihara H, Schlereth A, Cawthray GR, Encke B, Giavalisco P, Ivakov A, Arrivault S, Jost R, Krohn N, et al (2014) Low levels of ribosomal RNA partly account for the very high photosynthetic phosphorus-use efficiency of Proteaceae species. Plant Cell Environ 37: 1276–1298

Ticconi CA, Delatorre CA, Abel S (2001) Attenuation of phosphate starvation responses by phosphite in Arabidopsis. Plant Physiol 127: 963–972

Tran HT, Hurley BA, Plaxton WC (2010a) Feeding hungry plants: The role of purple acid phosphatases in phosphate nutrition. Plant Sci 179: 14–27

Tran HT, Plaxton WC (2008) Proteomic analysis of alterations in the secretome of Arabidopsis thaliana suspension cells subjected to nutritional phosphate deficiency. Proteomics 8: 4317–4326

Tran HT, Qian W, Hurley BA, She Y-M, Wang D, Plaxton WC (2010b) Biochemical and molecular characterization of AtPAP12 and AtPAP26: the predominant purple acid phosphatase isozymes secreted by phosphate-starved Arabidopsis thaliana. Plant Cell Environ 33: 1789–1803

Tripodi KE, Turner WL, Gennidakis S, Plaxton WC (2005) In vivo regulatory phosphorylation of novel phosphoenolpyruvate carboxylase isoforms in endosperm of developing castor oil seeds. Plant Physiol 139: 969–978

Tyanova S, Temu T, Cox J (2016) The MaxQuant computational platform for mass spectrometry-based shotgun proteomics. Nat Protoc 11: 2301–2319

Uhrig RG, Schläpfer P, Roschitzki B, Hirsch-Hoffmann M, Gruissem W (2019) Diurnal changes in concerted plant protein phosphorylation and acetylation in Arabidopsis organs and seedlings. Plant J 99: 176–194

Van der Does D, Boutrot F, Engelsdorf T, Rhodes J, McKenna JF, Vernhettes S, Koevoets I, Tintor N, Veerabagu M, Miedes E, et al (2017) The Arabidopsis leucine-rich repeat receptor kinase MIK2/LRR-KISS connects cell wall integrity sensing, root growth and response to abiotic and biotic stresses. PLoS Genet 13: e1006832

Veljanovski V, Vanderbeld B, Knowles VL, Snedden WA, Plaxton WC (2006) Biochemical and molecular characterization of AtPAP26, a vacuolar purple acid phosphatase up-regulated in phosphate-deprived Arabidopsis suspension cells and seedlings. Plant Physiol 142: 1282–1293

Veneklaas EJ, Lambers H, Bragg J, Finnegan PM, Lovelock CE, Plaxton WC, Price CA, Scheible W-R, Shane MW, White PJ, et al (2012) Opportunities for improving phosphorus-use efficiency in crop plants. New Phytol 195: 306–320

Versaw WK, Garcia LR (2017) Intracellular transport and compartmentation of phosphate in plants. Curr Opin Plant Biol 39: 25–30

Wakao S, Benning C (2005) Genome-wide analysis of glucose-6-phosphate dehydrogenases in Arabidopsis. Plant J 41: 243–256

Wang J, Lan P, Gao H, Zheng L, Li W, Schmidt W (2013) Expression changes of ribosomal proteins in phosphate- and iron-deficient Arabidopsis roots predict stress-specific alterations in ribosome composition. BMC Genomics 14: 783

Wang L, Lu S, Zhang Y, Li Z, Du X, Liu D (2014) Comparative genetic analysis of Arabidopsis purple acid phosphatases AtPAP10, AtPAP12, and AtPAP26 provides new insights into their roles in plant adaptation to phosphate deprivation. J Integr Plant Biol 56: 299–314

Wang T, Xing J, Liu Z, Zheng M, Yao Y, Hu Z, Peng H, Xin M, Zhou D, Ni Z (2019) Histone acetyltransferase GCN5-mediated regulation of long non-coding RNA At4 contributes to phosphate starvation response in Arabidopsis. J Exp Bot 70: 6337–6348

Wimalasekera R, Pejchar P, Holk A, Martinec J, Scherer GFE (2010) Plant phosphatidylcholine-hydrolyzing phospholipases C NPC3 and NPC4 with roles in root development and brassinolide signaling in Arabidopsis thaliana. Mol Plant 3: 610–625

Wiśniewski JR, Zougman A, Nagaraj N, Mann M (2009) Universal sample preparation method for proteome analysis. Nat Methods 6: 359–362

Yang J, Xie M-Y, Yang X-L, Liu B-H, Lin H-H (2019) Phosphoproteomic profiling reveals the importance of CK2, mapks and cdpks in response to phosphate starvation in rice. Plant Cell Physiol 60: 2785–2796

Yu B, Xu C, Benning C (2002) Arabidopsis disrupted in SQD2 encoding sulfolipid synthase is impaired in phosphate-limited growth. Proc Natl Acad Sci USA 99: 5732–5737

Zhang H, Liu D, Yang B, Liu W-Z, Mu B, Song H, Chen B, Li Y, Ren D, Deng H, et al (2020) Arabidopsis CPK6 positively regulates ABA signaling and drought tolerance through phosphorylating ABA-responsive element-binding factors. J Exp Bot 71: 188–203

Zhang W, Wang C, Qin C, Wood T, Olafsdottir G, Welti R, Wang X (2003) The oleate-stimulated phospholipase D, PLDdelta, and phosphatidic acid decrease H2O2-induced cell death in Arabidopsis. Plant Cell 15: 2285–2295

Zhang Z, Liao H, Lucas WJ (2014) Molecular mechanisms underlying phosphate sensing, signaling, and adaptation in plants. J Integr Plant Biol 56: 192–220

Zou J-J, Li X-D, Ratnasekera D, Wang C, Liu W-X, Song L-F, Zhang W-Z, Wu W-H (2015) Arabidopsis CALCIUM-DEPENDENT PROTEIN KINASE8 and CATALASE3 Function in Abscisic Acid-Mediated Signaling and H2O2 Homeostasis in Stomatal Guard Cells under Drought Stress. Plant Cell 27: 1445–1460

Zulawski M, Schulze G, Braginets R, Hartmann S, Schulze WX (2014) The Arabidopsis Kinome: phylogeny and evolutionary insights into functional diversification. BMC Genomics 15: 548

