## Supplemental Data 1 for "Phosphate and phosphite differentially impact the proteome and phosphoproteome of Arabidopsis suspension cell cultures"

**Supplemental Data XXX:**

**Legend: Core Protein Kinase Domain Motifs**

**RDLKPEN** = Catalytic Loop

**DFG** = Mg2+ Binding Loop

**TxY** = MPK Activation Loop

***SxY*** = Putative CDPK Activation Loop?

**APE** = End of P+1 loop

**Calcium Dependent Protein Kinases (CDPKs)**

**CDPK3 / CPK6 - AT2G17290**

MGNSCRGSFKDKIYEGNHSRPEENSKSTTTTVSSVHSPTTDQDFSKQNTNPALVIPVKEPIMRRNVDNQSYYVLGHKTPNIRDLYTLSRKLGQGQFGTTYLCTDIATGVDYACKSISKRKLISKEDVEDVRREIQIMHHLAGHKNIVTIKGAYEDPLYVHIVMELCAGGELFDRIIHRGHYSERKAAELTKIIVGVVEACHSLGVMH**RDLKPEN**FLLVNKDDDFSLKAI**DFG**LSVFFKPGQIFKDVVG***SPY***YV**APE**VLLKHYGPEADVWTAGVILYILLSGVPPFWAETQQGIFDAVLKGYIDFDTDPWPVISDSAKDLIRKMLCSSPSERLTAHEVLRHPWICENGVAPDRALDPAVLSRLKQFSAMNKLKKMALKVIAESLSEEEIAGLRAMFEAMDTDNSGAITFDELKAGLRRYGSTLKDTEIRDLMEAADVDNSGTIDYSEFIAATIHLNKLEREEHLVSAFQYFDKDGSGYITIDELQQSCIEHGMTDVFLEDIIKEVDQDNDGRIDYEEFVAMMQKGNAGVGRR**T**MKN**S**LNI**S**MRDV

**CDPK6 / CPK3 - AT4G23650**

MGHRHSKSKSSDPPPS**SSS**SSSGNVVHHVKPAGERRGSSGSGTVGSSGSGTGGSRSTTSTQQNGRILGRPMEEVRRTYEFGRELGRGQFGVTYLVTHKETKQQVACKSIPTRRLVHKDDIEDVRREVQIMHHLSGHRNIVDLKGAYEDRHSVNLIMELCEGGELFDRIISKGLYSERAAADLCRQMVMVVHSCHSMGVMH**RDLKPEN**FLFLSKDENSPLKAT**DFG**LSVFFKPGDKFKDLVG***SAY***YV**APE**VLKRNYGPEADIWSAGVILYILLSGVPPFWGENETGIFDAILQGQLDFSADPWPALSDGAKDLVRKMLKYDPKDRLTAAEVLNHPWIREDGEASDKPLDNAVLSRMKQFRAMNKLKKMALKVIAENLSEEEIIGLKEMFKSLDTDNNGIVTLEELRTGLPKLGSKISEAEIRQLMEAADMDGDGSIDYLEFISATMHMNRIEREDHLYTAFQFFDNDNSGYITMEELELAMKKYNMGDDKSIKEIIAEVDTDRDGKINYEEFVAMMKKGNPELVPNRRRM

**CDPK19 / CPK8 - AT5G19450**

MGNCCASPGSETGSKKGKPKIKSNPF**Y**SEAYTTNGSG**T**GFKLSVLKDPTGHDISLMYDLGREVGRGEFGITYLCTDIKTGEKYACKSISKKKLRTAVDIEDVRREVEIMKHMPRHPNIVSLKDAFEDDDAVHIVMELCEGGELFDRIVARGHYTERAAAAVMKTILEVVQICHKHGVMH**RDLKPEN**FLFANKKETSALKAI**DFG**LSVFFKPGEGFNEIVG***SPY***YM**APE**VLRRNYGPEVDIWSAGVILYILLCGVPPFWAETEQGVAQAIIRSVIDFKRDPWPRVSETAKDLVRKMLEPDPKKRLSAAQVLEHSWIQNAKKAPNVSLGETVKARLKQFSVMNKLKKRALRVIAEHLSVEEVAGIKEAFEMMDSKKTGKINLEELKFGLHKLGQQQIPDTDLQILMEAADVDGDGTLNYGEFVAVSVHLKKMANDEHLHKAFSFFDQNQSDYIEIEELREALNDEVDTNSEEVVAAIMQDVDTDKDGRISYEEFAAMMKAGTDWRKASRQYSRERFNSLSLKLMREGSLQLEGEN

**CDPK-like - AT1G49580**

MGGCTSKPSTSSGRPNPFAPGNDYPQIDDFAPDHPGKSPIPTPSAAKA**S**PFFPFY**T**P**S**PARHRRNKSRDVGGGGESK**S**LT**S**TPLRQLRRAFHPPSPAKHIRAALRRRKGKKEAALSGVTQLTTEVPQREEEEEVGLDKRFGFSKEFHSRVELGEEIGRGHFGYTCSAKFKKGELKGQVVAVKIIPKSKMTTAIAIEDVRREVKILQALSGHKNLVQFYDAFEDNANVYIAMELCEGGELLDRILARGGKYSENDAKPVIIQILNVVAFCHFQGVVH**RDLKPEN**FLYTSKEENSQLKAI**DFG**LSDFVRPDERLNDIVG***SAY***YV**APE**VLHRSYTTEADVWSIGVIAYILLCGSRPFWARTESGIFRAVLKADPSFDEPPWPFLSSDAKDFVKRLLFKDPRRRMSASQALMHPWIRAYNTDMNIPFDILIFRQMKAYLRSSSLRKAALRALSKTLIKDEILYLKTQFSLLAPNKDGLITMDTIRMALASNATEAMKESRIPEFLALLNGLQYRGMDFEEFCAAAINVHQHESLDCWEQSIRHAYELFDKNGNRAIVIEELASELGVGPSIPVHSVLHDWIRHTDGKLSFFGFVKLLHGVSVRASGKTTR

**Mitogen Activate Protein Kinases (MPKs)**

**MPK4 - AT4G01370**

MSAESCFGSSGDQSSSKGVATHGGSYVQYNVYGNLFEVSRKYVPPLRPIGRGAYGIVCAATNSETGEEVAIKKIGNAFDNIIDAKRTLREIKLLKHMDHENVIAVKDIIKPPQRENFNDVYIVYELMDTDLHQIIRSNQPLTDDHCRFFLYQLLRGLKYVHSANVLH**RDLKPSN**LLLNANCDLKLG**DFG**LARTKSETDFM**TEY**VVTRWYR**APE**LLLNCSEYTAAIDIWSVGCILGETMTREPLFPGKDYVHQLRLITELIGSPDDSSLGFLRSDNARRYVRQLPQYPRQNFAARFPNMSAGAVDLLEKMLVFDPSRRITVDEALCHPYLAPLHDINEEPVCVRPFNFDFEQPTLTEENIKELIYRETVKFNPQDSV

**MPK6 - AT2G43790**

MDGGSGQPAADTEMTEAPGGFPAAAPSPQMPGIENIPATLSHGGRFIQYNIFGNIFEVTAKYKPPIMPIGKGAYGIVCSAMNSETNESVAIKKIANAFDNKIDAKRTLREIKLLRHMDHENIVAIRDIIPPPLRNAFNDVYIAYELMDTDLHQIIRSNQALSEEHCQYFLYQILRGLKYIHSANVLH**RDLKPSN**LLLNANCDLKIC**DFG**LARVTSESDFM**TEY**VVTRWYR**APE**LLLNSSDYTAAIDVWSVGCIFMELMDRKPLFPGRDHVHQLRLLMELIGTPSEEELEFLNENAKRYIRQLPPYPRQSITDKFPTVHPLAIDLIEKMLTFDPRRRITVLDALAHPYLNSLHDISDEPECTIPFNFDFENHALSEEQMKELIYREALAFNPEYQQ

**MPK9 - AT3G18040**

MDPHKKVALETEFFTEYGEASRYQIQEVIGKGSYGVVASAIDTHSGEKVAIKKINDVFEHVSDATRILREIKLLRLLRHPDIVEIKHVMLPPSRREFRDIYVVFELMESDLHQVIKANDDLTPEHYQFFLYQLLRGLKFIHTANVFH**RDLKPKN**ILANSDCKLKIC**DFG**LARVSFNDAPSAIFW**TDY**VATRWYR**APE**LCGSFFSKYTPAIDIWSIGCIFAEMLTGKPLFPGKNVVHQLDIMTDLLGTPPPEAIARIRNEKARRYLGNMRRKPPVPFTHKFPHVDPLALRLLHRLLAFDPKDRPSAEEALADPYFYGLANVDREPSTQPIPKLEFEFERRKITKEDVRELIYREILEYHPQMLQEYLRGGEQTSFMYPSGVDRFKRQFAHLEENYGKGEKGSPLQRQHASLPRERVPAPKKENGSHNHDIENRSIASLVTTLESPPTSQHEGSDYRNGTSQTGYSARSLLKSASISASKCIGMKPRNKSEYGESNNDTVDALSQKVAALHT

**MPK15 - AT1G73670**

MGGGGNLVDGVRRWLFFQRRPSSSSSSNNHDQIQNPPTVSNPNDDEDLKKLTDPSKLRQIKVQQRNHLPMEKKGIPNAEFFTEYGEANRYQIQEVVGKGSYGVVGSAIDTHTGERVAIKKINDVFDHISDATRILREIKLLRLLLHPDVVEIKHIMLPPSRREFRDVYVVFELMESDLHQVIKANDDLTPEHHQFFLYQLLRGLKYVHAANVFH**RDLKPKN**ILANADCKLKIC**DFG**LARV**S**FNDAPTAIFW**TDY**VA**T**RWYR**APE**LCGSFFSKYTPAIDIWSVGCIFAEMLLGKPLFPGKNVVHQLDIMTDFLGTPPPEAISKIRNDKARRYLGNMRKKQPVPFSKKFPKADPSALRLLERLIAFDPKDRPSAEEALADPYFNGLSSKVREPSTQPISKLEFEFERKKLTKDDIRELIYREILEYHPQMLEEYLRGGNQLSFMYPSGVDRFRRQFAHLEENQGPGGRSNALQRQHASLPRERVPASKNETVEERSNDIERRTTAAVASTLDSPKASQQAEGTENGGGGGYSARNLMKSSSISGSKCIGVQSKTNIEDSIVEEQDETVAVKVASLHNS
