## Supplementary figures and images for "Phosphate and phosphite differentially impact the proteome and phosphoproteome of Arabidopsis suspension cell cultures"

### Figure S1

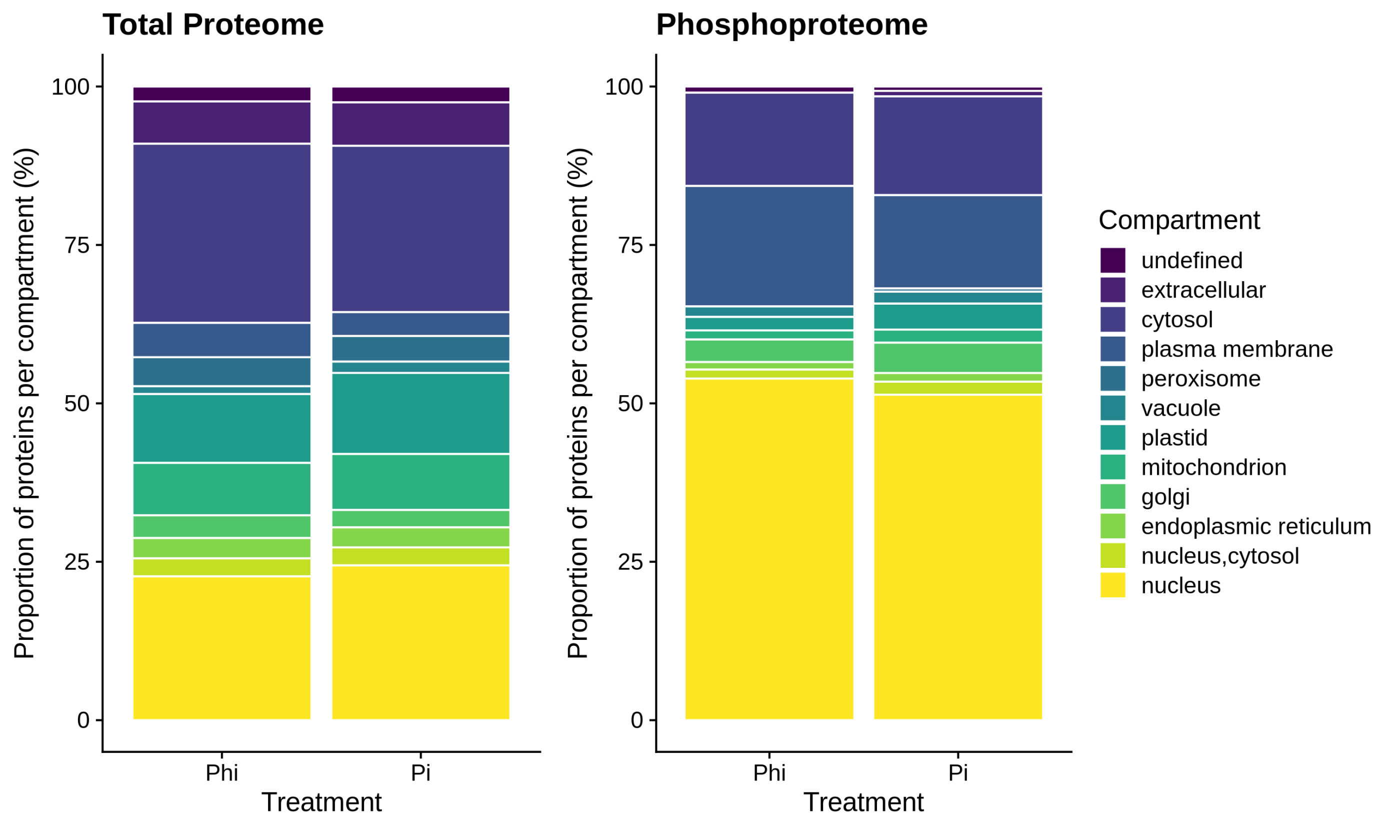

### Figure S2

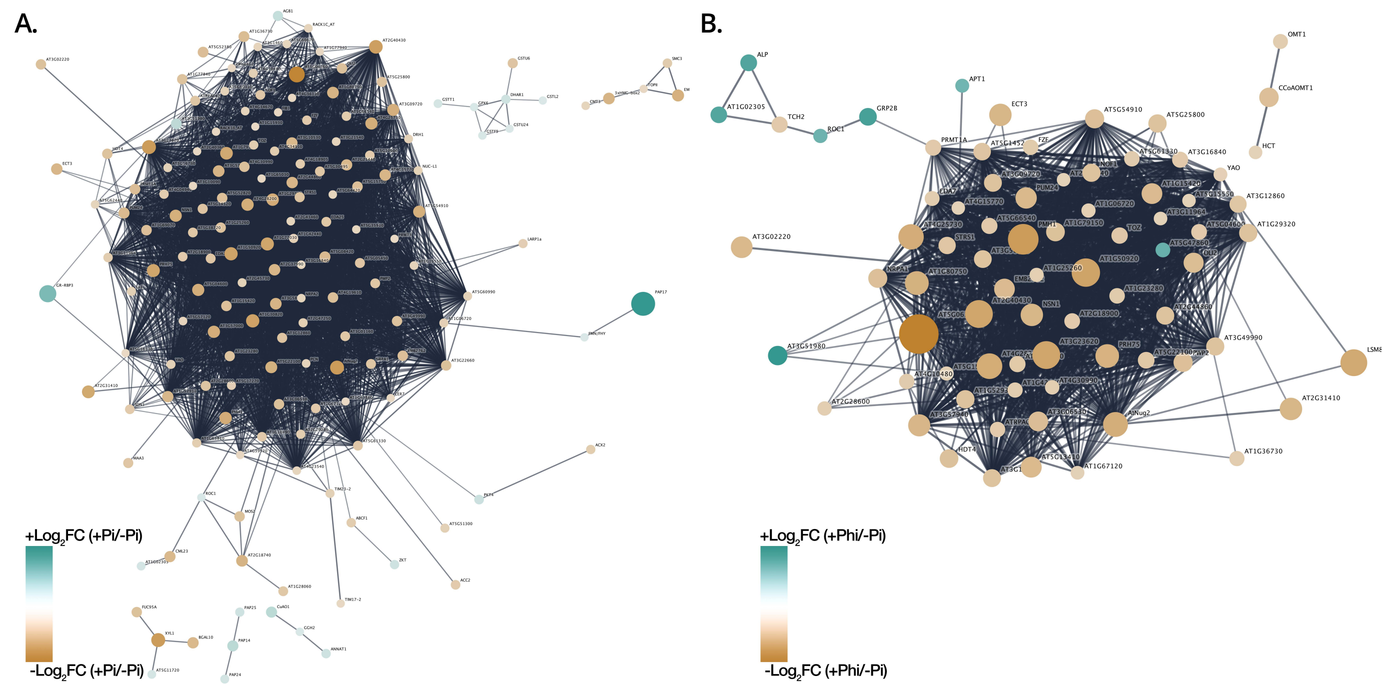
